# Vertical stratification drives additive prokaryotic diversity in beech forest floors, while site conditions shape boundary layers

**DOI:** 10.64898/2026.06.02.729534

**Authors:** Sebastian Bibinger, Jörg Niederberger, Friederike Lang, Michael Schloter, Stefanie Schulz

## Abstract

The forest floor is a key interface regulating carbon and nutrient processing and transfer to the mineral soil, yet it is threatened by climate-change-driven reductions in organic layer mass. How its prokaryotic microbiome is structured across the fine-scale vertical gradient from fresh litter to mineral topsoil remains poorly resolved. We characterised prokaryotic communities across eight sequential layers spanning fresh litter, organic layers at different stages of decomposition, and mineral topsoil in three temperate beech forests using 16S rRNA gene amplicon sequencing. The decomposition stage was the dominant driver of community assembly, resulting in a strongly vertically stratified prokaryotic microbiome. While alpha diversity peaked within the fragmented litter layers, the overall high diversity of the forest floor was primarily an additive effect of vertical stratification. Site-specific effects were pronounced in the litter layer and the mineral topsoil, but diminished in the intermediate organic layers, where high local heterogeneity masked between-site differences. Redundancy analysis further showed that the environmental drivers of community structure shifted with depth, from litter quality in the upper horizons to mineral-associated properties in the mineral topsoil. At the same time, predicted 16S rRNA gene copy numbers indicate that the humified layers harbour the highest abundance of oligotrophic life strategists in the profile. By resolving the fine-scale vertical structure of the forest floor prokaryotic microbiome, our results provide a baseline for predicting how climate-change induced loss of organic layer mass threatens layer-specific communities, particularly the oligotrophic taxa in the humified layers, with consequences for the carbon turnover, tree nutrition, and nutrient cycling functions they mediate.

## Introduction

The forest floor (FF) is the interface between the mineral soil and the atmosphere in forest ecosystems. It forms through the accumulation and decomposition of organic matter, primarily plant litter. The FF is central to biogeochemical processes in forests, representing a substantial carbon stock and the primary pathway for the transport of nutrients and organic matter to the mineral soil (Binkley & Fisher, 2020; Lang et al., 2025; Ponge, 2013). Additionally, it fulfils crucial functions, like providing a medium for tree seedling establishment, retaining water, and preventing soil erosion (Floriancic et al., 2022; Sayer, 2006; Zhang et al., 2022; F. Zhu & Cheng, 2022; J. Zhu et al., 2023). Especially under nutrient poor conditions, the FF contributes to tree nutrition in addition to the mineral soil (Gao et al., 2021; Lang et al., 2016; Sayer et al., 2024). Finally, the FF hosts a diverse community of fauna, fungi, and bacteria. While fungi are thought to be the main drivers of leaf litter degradation due to their prevalent capability to degrade lignin and cellulose, the role of bacteria is increasingly recognised (Bani et al., 2018; Glassman et al., 2018; López-Mondéjar et al., 2016; Štursová et al., 2012). Both groups are of paramount importance for the (re-) cycling of organic matter in forest ecosystems (Baldrian, 2017; Lindahl et al., 2007). The dominant tree species is the most critical factor shaping the FF properties and associated microbiomes, since it defines the litter quantity and quality (Laganière et al., 2010; Lladó et al., 2018; Urbanová et al., 2015). However, the FF is further shaped by abiotic factors, such as precipitation, temperature, and, notably, the underlying soil parent material through both top-down and bottom-up pathways (García-Palacios et al., 2016; Kaiser et al., 2012). For instance, calcium brought into the FF through leaf litter can slow decomposition by protecting organic matter (Rowley et al., 2018). Iron and aluminium, which can be actively transported from the mineral soil into the FF by ectomycorrhizal fungi or bioturbation, regulate the pH in acidic environments and facilitate organic matter transformation (Clarholm & Skyllberg, 2013).

Additionally, being composed of litter in various stages of decomposition, the FF is heavily vertically stratified. According to the German soil taxonomy, it can be separated into an Ol layer (undecomposed leaves, <10% organic fine matter, OFM), a partially degraded Olf (<35% OFM) and Ohf (<70% OFM) layers, and a Oh layer (>70% OFM), which consists mostly of organic fine matter (Wachendorf et al., 2023). Consequently, strong changes in physicochemical properties are observed with increasing depth in the FF. This shift is characterised by the progressive transformation of organic matter, starting with the leaching of labile compounds such as sugars and amino acids, followed by the subsequent decomposition of major structural polymers such as cellulose and lignin, accompanied by a decrease in C:N and C:P ratios (Šnajdr et al., 2011). As labile compounds are rapidly depleted, recalcitrant high-molecular-weight aromatic residues and microbial necromass accumulate, leading to the formation of humic substances (Krishna & Mohan, 2017; Kulikova & Perminova, 2021; Prescott & Vesterdal, 2021). At the same time, an increasing abundance of minerals brought in from the mineral soil through bioturbation can be expected (Clarholm & Skyllberg, 2013).

Vertical stratification is a well-known driver of microbial community composition in soil environments (Hao et al., 2021; Naylor et al., 2022; Zhao et al., 2021), and due to the strong physicochemical gradient, this is particularly true in forest soils (Baldrian, 2017; De Araujo Pereira et al., 2017; Jiao et al., 2018). For example, the fungal-to-bacterial ratio decreases with depth. Saprotrophic fungi, which dominate in the litter layer, are replaced by bacteria in the lower organic horizon and the mineral soil (Baldrian et al., 2012). Prokaryotic communities shift from dominance of copiotrophic to oligotrophic groups, and their cross-kingdom interactions are strengthened in the mineral soil compared to the organic horizon (López-Mondéjar et al., 2015; Mundra et al., 2021). Consequentially, the functional potential regarding litter degradation also changes with depth, adapting to the shifting substrate quality (Baldrian et al., 2012; Šnajdr et al., 2008; Uroz et al., 2013).

To disentangle how prokaryotic communities are structured by the progressive degradation of litter versus the increasing bottom-up influence of the mineral soil, a fine-scale vertical resolution at different sites is required. However, in previous research on FF stratification, this fine-scale resolution approach is underrepresented. While studies using time-resolved samplings are abundant and can resolve microbiome succession during leaf litter decomposition (Buresova et al., 2019; Gołębiewski et al., 2019; Maillard et al., 2024; Min et al., 2023; Schroeter et al., 2022), the turnover of the forest floor in temperate forests exceeds most study timeframes (Adams et al., 2019; Currie, 1999; Raza et al., 2023).

Resolving these spatial dynamics is urgent, as climate change-induced forest change puts forest floors at risk. Climate change and the following increased FF turnover may lead to a decrease in FF mass and the loss of specific ecological niches with their associated microbiome (Lang et al., 2025).

Thus, this study aims to characterise the vertical stratification of the prokaryotic microbiome within the forest floor organic layers and topsoil horizon. We sampled these layers at fine vertical resolution and performed 16S rRNA gene amplicon sequencing across three beech-dominated German forests with similar climates. We further analysed a range of physicochemical parameters to determine how site-specific differences influence the prokaryotic community and nutrient availability across different layers of the forest floor and mineral topsoil.

We hypothesise that the state of litter decomposition is the primary predictor of prokaryotic community structure. Consequently, microbiome diversity, composition, and predicted growth rates will exhibit strong vertical stratification, which is independent of site. However, we expect site-specific physicochemical parameters to drive species identity, particularly at the upper and lower boundaries of the forest floor, with litter quality (top-down) and interactions with mineral soil (bottom-up) mediating these effects.

## Material and Methods

### Soil Sampling

This study focused on three European beech (*Fagus sylvatica L.*) dominated forests in southern Germany, Bad Brückenau (BBR), Kandel (KAN), and Mitterfels (MIT). The exact location and properties are listed in Table 1. Forest floor samples were taken in October of 2022 after litterfall but before snowfall.

**Table 1:** Environmental characteristics and site parameters of the surveyed beech forests. Mean annual temperature (MAT) and mean annual precipitation (MAP) refer to the period 1991–2020 (Deutscher Wetterdienst, 2026). Soil type refers to the World Reference Base (WRB) 3rd edition 2015 (BBR and MIT) and 4th edition 2022 (KAN) (IUSS Working Group WRB, 2015, 2022).

| Site | Location | Latitude | Longitude | Elevation (a.s.l.) | MAT | MAP | Dominant humus form | Soil Type (World Reference Base) | Bedrock Type |
| --- | --- | --- | --- | --- | --- | --- | --- | --- | --- |
| Bad<br>Brücke<br>nau<br>(BBR) | Rhön<br>Mountai<br>ns | 50.35<br>2551 | 9.927<br>604 | 809 m | 6.8 °C | 1110<br>mm | Typical<br>F-Mull | Eutric<br>Cambi<br>sol | Tertiär<br>y<br>Volcan<br>ic<br>Basalt |
| Kandel<br>(KAN) | Black<br>Forest | 48.06<br>8709 | 8.026<br>324 | 1163<br>m | 6.3 °C | 1769<br>mm | Typical<br>Moder | Dystric<br>Skeleti<br>c<br>Rhodic<br>Cambi<br>sol | Parag<br>neiss |
| Mitterfel<br>s<br>(MIT) | Bavaria<br>n Forest | 48.97<br>635 | 12.87<br>69 | 990 m | 6.0 °C | 1448<br>mm | Typical<br>Moder | Dystric<br>Cambi<br>sol | Parag<br>neiss |

Five representative quantitative pits (q-pits, 0.4 × 0.4 m) were excavated at each site. To ensure representativeness, sampling locations were selected in the central crown region of adult beech trees, at least 2 m from the stem and never directly downslope, to avoid the influence of stemflow. Areas with understory vegetation or deadwood were excluded.

The forest floor was classified and then sampled carefully from top to bottom. Samples were taken in up to five layers (Ol0, Ol1, Olf, Ohf, Oh) in fine vertical resolution. Ol0 represents the freshly fallen leaves, while Ol1 represents litter with residence time longer than approximately two weeks. Olf, Ohf, and Oh were classified according to the updated version of the German humus form systematics (Wachendorf et al., 2023). Two additional samples per pit were taken from the underlying mineral soil at depths of 0–5 cm (A5) and 5–10 cm (A10).

In the field, samples from the Oh and the A horizon were homogenised by sieving (2 mm mesh), after which subsamples were flash-frozen on dry ice. Due to their high variability, layers Ol0, Ol1, Olf, and Ohf were collected as larger bulk samples (approx. 50 ml) and frozen as a whole. All samples were subsequently stored at -80 °C. Before DNA extraction, samples from layers Ol0-Ohf were ground in liquid nitrogen to ensure homogenization.

### Soil Chemical Analysis

Soil and forest floor pH was determined in air-dried samples using a 1 M KCl solution at a soil-to-solution ratio of 1:5. Measurements were performed using a Metrohm CH/789 Robotic Sample Processor XL (Metrohm, Switzerland) equipped with a Micro El. Cone 16 WOC electrode.

Total carbon (C) and nitrogen (N) concentrations were quantified by dry combustion at 1100 °C using an EuroEA elemental analyser (HEKAtech, Germany).

Citrate-extractable phosphorus (P) fractions were analysed as described before (Gutachterausschuss Forstliche Analytik, 2014). Briefly, 5 g of soil or 2.5 g of forest floor material were extracted with 50 ml of 1% citric acid monohydrate, shaken for 2 h, and allowed to settle for 12–18 h. Total P in the citrate extract was determined using an Agilent 5800 ICP-OES (Agilent Technologies, USA).

Exchangeable cations were extracted using the NH_4_OAc method at pH 7, as described before (Lavkulich, 1981). Element concentrations were measured using a Vista-Pro CCD Simultaneous ICP-OES (Varian, Germany).

### DNA Extraction and Sequencing Library Preparation

Approximately 0.3 grams of soil were used per sample for DNA extraction. Bead beating was performed using a Precellys 24 tissue homogeniser (Bertin Instruments, France). DNA was extracted using the NucleoSpin Soil kit (Macherey-Nagel, Germany) with buffer SL2 and enhancer SX, following the manufacturer’s instructions, and eluted in 50 µl of water. The quality of the extracted DNA was assessed using a NanoDrop 1000 spectrophotometer (ThermoFisher Scientific, USA), and the quantity was measured using the Quant-iT PicoGreen dsDNA Assay Kit (Thermo Fisher Scientific, USA) with a Spark® Multimode Microplate Reader (Tecan Trading AG, Switzerland). Per round of extraction, one extraction control without soil was processed.

The V4 region of the 16S rRNA gene was amplified using the primer pair 515FB-806RB (Apprill et al., 2015; Parada et al., 2016). PCR amplification of the target region was performed in 25 µl reactions containing 5 ng of template DNA, 12.5 µl of NEBNext high-fidelity polymerase, 0.3 pmol of each primer, and 2.5 µl of 3% BSA. The thermal cycling profile consisted of an initial denaturation at 98°C for 1 min, followed by 25 cycles of 98°C for 10 s, 55°C for 30 s, and 72°C for 30 s, with a final extension at 72°C for 5 min. Successful amplification was confirmed via 1% (w/v) agarose gel electrophoresis. To control for contamination, a no-template control, in which the template DNA was replaced by nuclease-free water, was included with each set of PCR reactions. The negative controls were processed and sequenced alongside all other samples. Amplicons were then purified using MagSI NGSprep Plus Beads (Magtivio, Netherlands) at a 0.8:1 ratio, and the final DNA concentration was quantified with the Quant-iT PicoGreen dsDNA Assay Kit.

Sequencing indices were attached to the 16S via a second PCR using the Nextera XT Index Kit v2 (Illumina, Inc., USA). Each 25 µl indexing reaction contained 10 ng of template DNA, 12.5 µl of NEBNext high-fidelity polymerase, and 2.5 µl of each indexing primer. The thermal profile was an initial 30 s at 98°C, followed by 8 cycles of 98°C for 10 s, 55°C for 30 s, and 72°C for 30 s, with a final extension at 72°C for 5 min. The indexed products were purified using MagSI NGSprep Plus Beads and quantified by capillary electrophoresis on a 5200 Fragment Analyser (Agilent, USA). Finally, libraries were pooled in equimolar concentrations (4 nM, 20% PhiX) and sequenced on an Illumina MiSeq platform with the Reagent Kit v3.

### Sequence Data Preprocessing

Raw, demultiplexed 16S sequences were processed by first removing sequencing adapters with *CutAdapt* v3.5 (Martin, 2011). Subsequent read processing was performed using the *DADA2* pipeline v.1.34.0 (Callahan et al., 2016) within *R* v4.5.0 (R Core Team, 2025) and RStudio v2023.09.1 (Posit team, 2025). Forward and reverse reads were trimmed to 260 bp and 200 bp, respectively, and quality-filtered using maxEE = (2,2). The resulting ASV sequences were taxonomically classified against the *SILVA* database v138.1 (Quast et al., 2013). Reads classified to mitochondrial or chloroplast origins were removed (3.66% of total reads in 1244 ASVs). Furthermore, potential contaminants were then identified and removed (0.59% of total reads in 104 ASVs) using the *decontam* package v1.22.0 (Davis et al., 2018). After assessing sequencing depth with rarefaction curves using the vegan package v.2.6.10 (Oksanen et al., 2025), samples were rarefied prior to alpha diversity calculations and otherwise normalised using cumulative sum scaling (CSS) with the *metagMisc* package v.0.5.0 (Mikryukov, 2025). A phylogenetic tree was computed using *phangorn* v.2.12.1 (Schliep, 2011) and *IQ-TREE* 2 v.2.4.0 (Minh et al., 2020). ASVs with a total read count less than 10 across all samples were excluded from further analysis. Detailed read count information is available in Supplementary Table S1.

### Statistical Analysis

Statistical analysis was performed using *R* v4.5.0 in RStudio 2023.09.1 with *phyloseq* v.1.52.0 (McMurdie & Holmes, 2013). Unless specified differently, analyses were performed using the package *vegan* v.2.6.10 (Oksanen et al., 2025). Data were visualised using *ggplot* v.3.5.2 (Wickham, 2016), *ggokabeito* v.0.1.0 (Barrett, 2022), *viridis* v.0.6.5 (Garnier et al., 2024), *ComplexHeatmap* v.2.24.1 (Gu, 2022), and *ggpubr* v.0.6.1 (Kassambara, 2026).

One replicate q-pit from site KAN was excluded from subsequent analyses as a significant compositional outlier. Its group centroid exhibited a distance of 0.48 to the global centroid in PCoA space, which was 1.65 times greater than that of the next most distant group (Supplementary Figure S1).

Alpha diversity indices were calculated using the package *phyloseq* on samples rarefied to the minimum library size. Differences in alpha diversity (observed ASV and phylum richness) were analysed using Linear mixed-effects models (LMM) using the package *lme4* v.1.1.37 (Bates et al., 2015). Site and soil layer were specified as fixed effects (including their interaction), while the sampling pit was included as a random intercept to account for the nested sampling design. Richness data were log-transformed to meet model assumptions, which were verified via Q–Q plots, the Shapiro–Wilk test for normality, and Levene’s test from the package *car* v.3.1.5 (Fox & Weisberg, 2019) for homogeneity of variance. The significance of fixed effects was assessed using Type III analysis of variance (ANOVA) with Satterthwaite’s method for approximating degrees of freedom using the package *lmerTest* v.3.1.3 (Kuznetsova et al., 2017). Estimated marginal means and pairwise post-hoc comparisons (Tukey-adjusted) were computed using the package *emmeans* v.2.0.1 (Lenth et al., 2026). Since the Oh layer was not present at BBR, pairwise comparisons among sites were averaged over the six layers present at all sites.

Weighted UniFrac distances between samples were computed using the “distance” function from *phyloseq*. To test for significant differences in community composition between sites and layers, a Permutational multivariate analysis of variance (PERMANOVA) with 999 permutations was performed on weighted UniFrac distances with 999 permutations. The permutation was constrained by sampling q-pit. Homogeneity of variances was confirmed beforehand. Non-metric multidimensional scaling (NMDS) was used to visualise differences in community composition.

16S rRNA gene copy number/genome (16S GCN) was predicted from ASV sequences using ANNA16 software v.1.1.0 (Miao et al., 2024). The abundance-weighted average 16S GCN was then calculated per sample. Statistical testing was performed using linear mixed models, as for alpha diversity measures.

Indicator genera per site and horizon were identified through calculating the *IndVal.g* index using *indicspecies* v.1.8.0 (De Cáceres & Legendre, 2009). The analysis was carried out on a dataset aggregated to genus level and filtered for the core microbiome. Layer-specific core microbiomes were computed using the package *microbiome* v.1.30.0 (Lahti & Shetty, 2017) with a detection cutoff of 0.3% and a prevalence of 80% per site.

The influence of soil physicochemical parameters on prokaryotic community structure was evaluated using distance-based redundancy analysis (dbRDA) on weighted UniFrac distances, using the *vegan* package. A global dbRDA model was fitted across all soil layers. Forward selection of explanatory variables was performed using ordiR2step (999 permutations) based on the permutation p-value (α = 0.05) and the adjusted R² value. Layer-specific dbRDA models were fitted separately to identify layer-specific drivers. To avoid overfitting given the limited sample size per layer, variables were pre-selected per layer based on significant variation between sites (ANOVA) and significant explanatory power for community structure (PERMANOVA on weighted UniFrac distances, 999 permutations). Associated p-values were adjusted using the Benjamini-Hochberg method and variables with q < 0.05 in both tests were retained. Subsequently, variables with a variance inflation factor > 10 were identified through the package *usdm* v.2.1.7 (Naimi et al., 2014) and removed. Retained environmental factors were used for dbRDA on weighted UniFrac distances. The significance of the overall and layer-specific models, as well as individual terms, was assessed using ANOVA-like permutation tests (999 permutations).

### Use of Artificial Intelligence tools

The large-language models Gemini Pro (Google LLC) and Claude Opus (Anthropic, PBC) were used for language editing of the manuscript text and for assistance with formatting and debugging of R analysis scripts. No AI tools were used to generate results or their interpretation. All AI-generated content and edits were reviewed and adapted by the authors, who take full responsibility for the content of the publication.

## Results

### Alpha and beta diversity patterns follow FF stratification

Following bioinformatic pre-processing, an average of 44,684 high-quality reads per sample were obtained. Rarefaction analysis confirmed that this sequencing depth was sufficient to capture the microbial diversity across the samples (Supplementary Figure S2). The final dataset comprised 11,844 unique amplicon sequence variants (ASVs), 11,742 of which were assigned to Bacteria and 102 to Archaea.

Soil layer was the dominant factor influencing ASV richness (Figure 1A; LMM F = 28.6, p < 0.001), while site played a secondary but still significant role (F = 6.02, p = 0.017).

**Figure 1:**
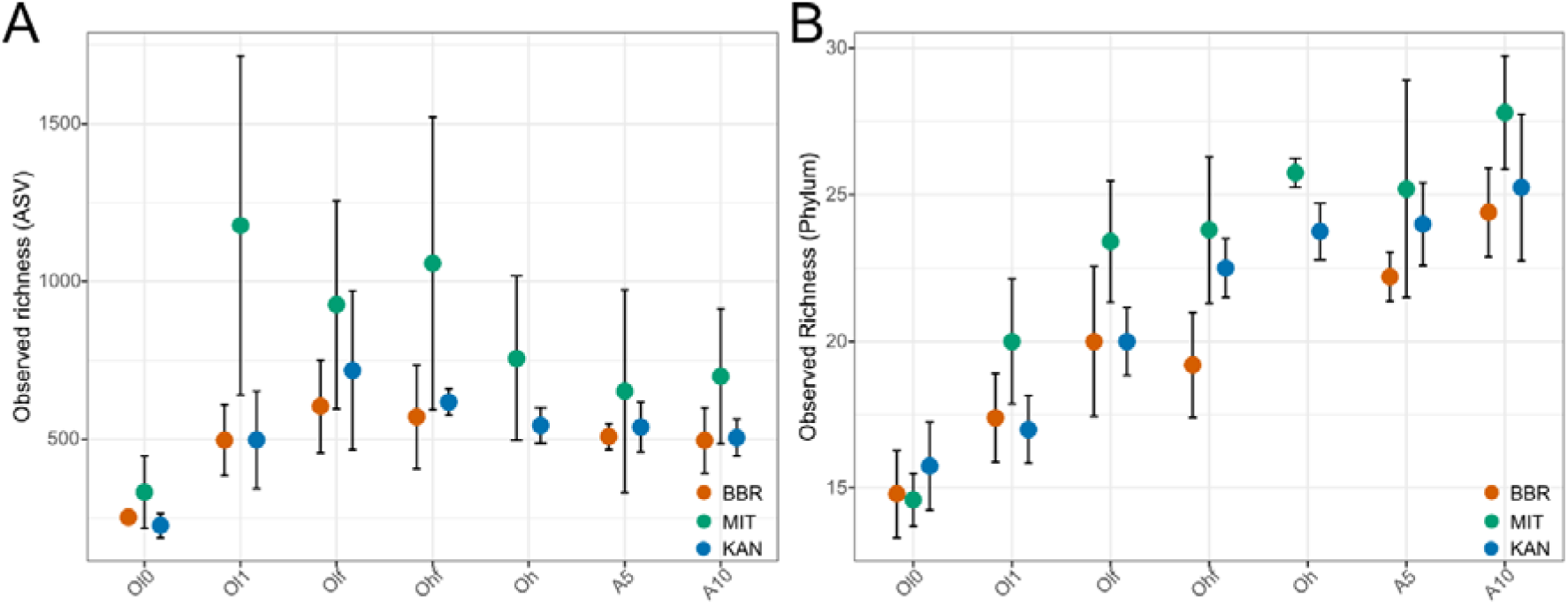
Prokaryotic microbiome alpha diversity across the forest soil profile. Observed richness of the prokaryotic community across forest floor layers at the ASV (A) and phylum level (B), shown for three forest sites: Bad Brückenau (BBR), Mitterfels (MIT), and Kandel (KAN). Data points represent group means with whiskers indicating standard deviation (n = 5 for BBR and MIT; n = 4 for KAN; the Oh layer was sampled only at MIT and KAN). Richness was estimated on samples rarefied to the minimum library size. Differences in richness were assessed using linear mixed-effects models with site, layer, and their interaction as fixed effects and sampling pit as a random factor; richness data were log-transformed prior to analysis. Full statistical results are provided in Supplementary Tables S2–S4.

Observed richness was lowest in the Ol0 layer (273 ± 81 ASVs, p < 0.001 for all pairwise comparisons) and increased to its apparent peak in the Olf (751 ± 274) and Ohf layers (758 ± 359). Following this peak, richness decreased in the Oh layer (649 ± 209) and into the underlying mineral soil (A1: 568 ± 195; A2: 570 ± 168). However, these differences among the deeper layers were not statistically significant after correction for multiple comparisons, likely caused by high within-layer variability, particularly at site Mitterfels. Overall, Mitterfels exhibited significantly higher richness than both Bad Brückenau (p = 0.018) and Kandel (p = 0.040).

Observed phylum richness exhibited a different pattern (Figure 1B). Soil layer was again the dominant factor (LMM F = 70.4, p < 0.001) over site (F = 10.2, p = 0.003). Their interaction was marginally non-significant (F = 1.84, p = 0.066). In contrast to the ASV-level analysis, phylum richness showed a monotonic increase with depth from the Ol0 to the mineral soil (p < 0.01 for all comparisons but Olf–Ohf and A1–A2). Among sites, only Bad Brückenau exhibited significantly lower phylum richness than Mitterfels (p = 0.002).

Complete ANOVA tables and pairwise post-hoc comparisons are provided in Supplementary Tables S2–S4. Results for the Shannon diversity and Pielou’s evenness show comparable trends (Supplementary Figure S3).

The soil layer was also the strongest driver of the prokaryotic microbiome composition (PERMANOVA on weighted UniFrac distances: R² = 0.82, p < 0.001) while site alone played a smaller role (R² = 0.04, p < 0.001). Still, a significant interaction between layer and site was observed (R² = 0.07, p < 0.001). Accordingly, non-metric Multidimensional Scaling (NMDS) revealed clear clustering by layer and a continuous shift in community composition from the fresh litter layers (Ol0) down to the lower mineral soil layer (A10) (Figure 2A, Supplementary Table S5).

**Figure 2:**
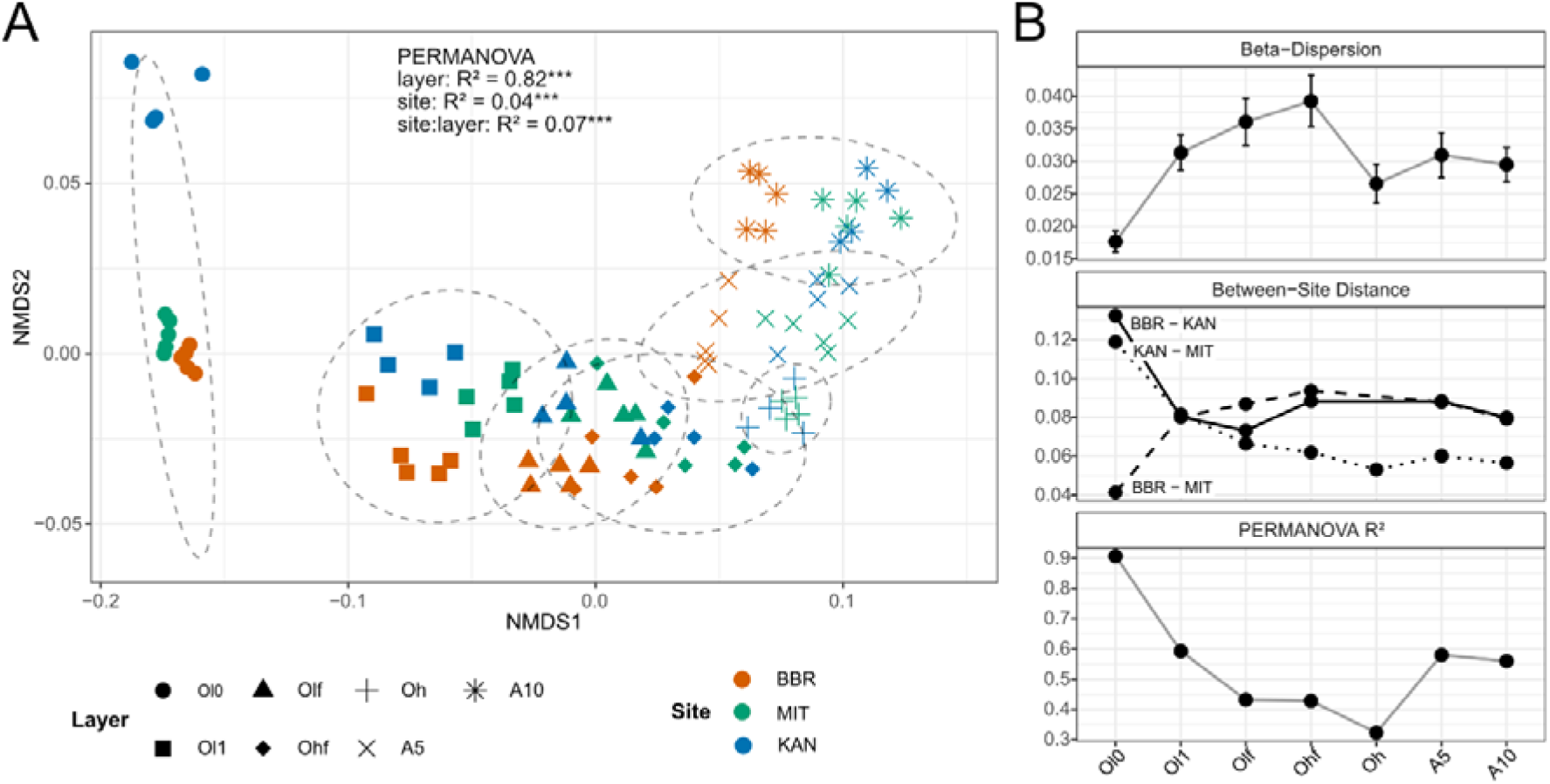
Prokaryotic microbiome beta diversity across the soil layers. (A) Non-metric multidimensional scaling (NMDS) ordination based on weighted UniFrac distances. Colours correspond to the site, while shapes represent the layer. Ellipses represent 95% confidence clusters. The NMDS stress value was 0.042. (B) Changes in beta-diversity metrics along the soil profile. Displayed are mean beta-dispersion (top), mean distance between sites split by comparison (middle), and layer-wise PERMANOVA R^2^ for factor site (bottom). Error bars indicate standard error.

Beta-diversity metrics varied substantially across the soil profile (Figure 2B). The explanatory power of the site was highest in the litter layer Ol0 (R² = 0.91), where beta-dispersion was lowest (0.018 ± 0.002) and between-site distances varied strongly. Kandel showed the highest pairwise distances across the entire profile (0.132 and 0.119 to Bad Brückenau and Mitterfels, respectively), while Bad Brückenau and Mitterfels were notably more similar (0.041). Subsequently, pairwise distances converged rapidly in layer Ol1 (∼0.080 for all comparisons) and then diverged again. While Mitterfels and Kandel prokaryotic communities became more similar with depth in the FF (to 0.053 in the Oh), the community from Bad Brückenau remained consistently relatively distinct from the other two sites (0.079–0.094). Beta-dispersion increased continuously up to layer Ohf (0.039) and subsequently dropped again to an intermediate level (0.029 ± 0.002). Together, these trends drove a decline in the site effect to its minimum in the Oh layer (R² = 0.32), followed by a recovery in the mineral soil layers, where the site effect stabilised (R² = 0.58 and 0.56 for A5 and A10, respectively). We repeated the analysis using only the two complete datasets (Kandel and Mitterfels) to avoid an unbalanced design for layer Oh. In the balanced dataset, the site effect reached its minimum in the Ohf layer (R² = 0.09), followed by a recovery in the Oh layer (R² = 0.32) driven by a decrease in beta-dispersion (Supplementary Figure S4).

### Taxonomic composition and lifestyle strategy follow FF stratification

At the phylum level, the compositional trends of the prokaryotic community across the sampled layers were largely identical among sites (Figure 3A). Although the highly abundant phyla *Pseudomonadota* (across sites and layers relative abundance = 35.8%), *Actinomycetota* (18.0%), and *Acidobacteriota* (12.6%) were ubiquitous across the entire surveyed soil profile, their relative abundances changed with depth. *Pseudomonadota* and *Bacteroidota* dominated the litter layers (e.g., Ol0: 58.2% and 27.9% respectively). Their relative abundance progressively declined towards the mineral soil (A10: 24.2% and 0.5%). In contrast, *Acidobacteriota* and *Chloroflexota* exhibited the reverse pattern. While they were rare in the litter horizons (Ol0: 0.6% and 0.1%), they became progressively more abundant with depth, peaking in the mineral soil layers (e.g., A10: 20.9% and 13.7%). A similar trend was observed for *Verrucomicrobiota*, *Thermoproteota*, and Candidatus *Eremiobacterota*. *Actinomycetota* and *Planctomycetota* exhibited a unimodal abundance pattern across the soil profile. Their relative abundance peaked in the organic layers Olf to Oh (Ohf values: 23.7% and 11.4%) before decreasing again in the mineral soil. Furthermore, some phyla were found exclusively in one layer: *Deinococcota* was restricted to the litter layers, whereas *Nitrospirota*, *Methylomirabilota*, NB1-j, and MBNT15 were found exclusively in the mineral soil. Complete phylum-level abundances layer and the most abundant genera per layer are provided in Supplementary Table S6 and Figure S5, respectively.

**Figure 3:**
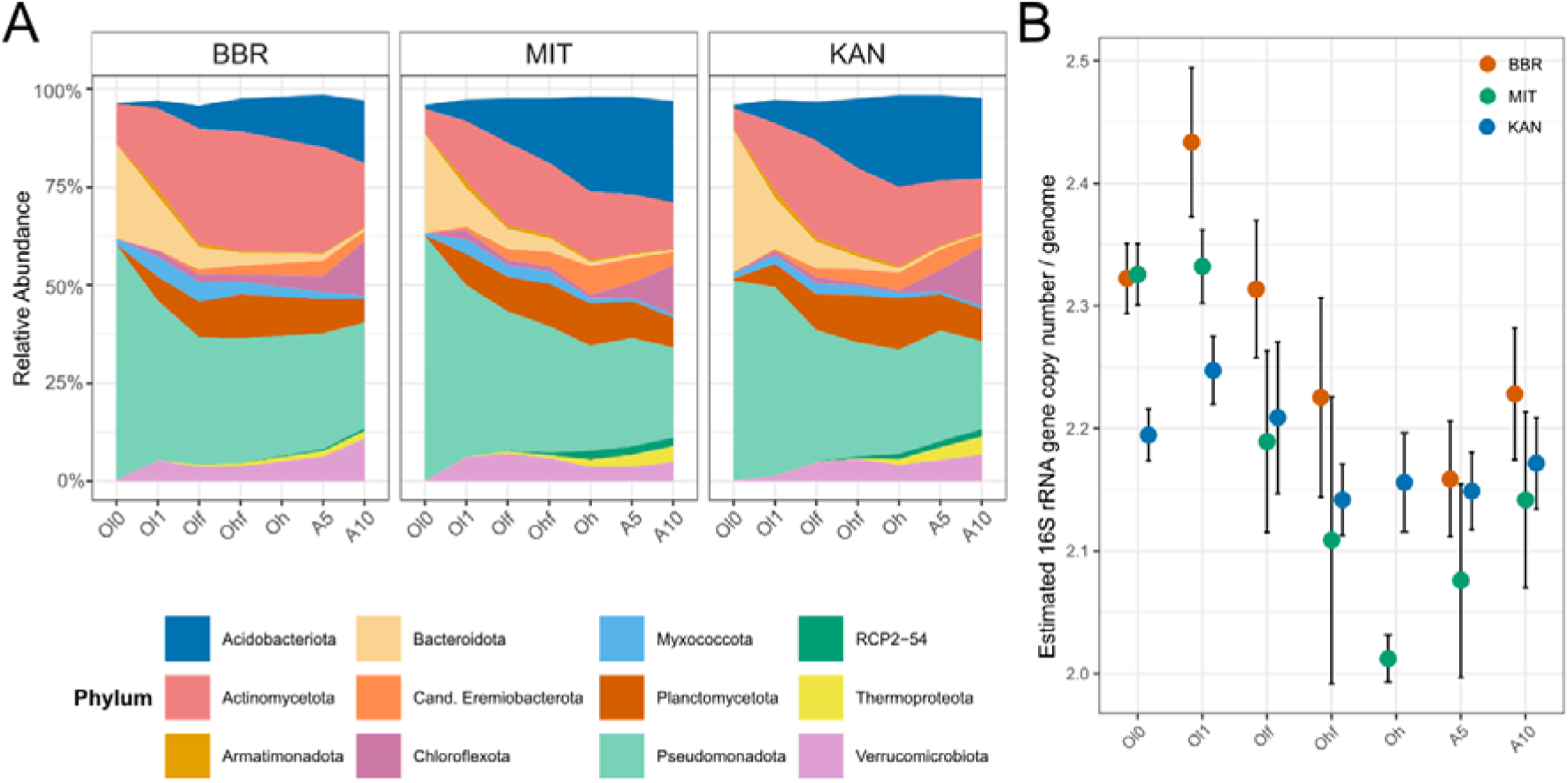
Microbial community structure across forest soil layers. (A) Stacked area plots showing the mean relative abundance of dominant bacterial phyla across the layers, faceted by site. Shown are the top 12 Phyla by abundance, which covered 97.3 % of all reads. (B) Average predicted, abundance-weighted, 16S gene copy number (GCN) per soil layer and site. A higher average 16S GCN indicates a higher presence of fast-growing taxa. Whiskers indicate standard deviation.

We estimated the average 16S rRNA gene copy number/genome (16S GCN) to assess how the prokaryotic average growth lifestyle changes along the profile (Figure 3B). A higher 16S GCN indicates a greater relative abundance of fast-growing bacteria. Overall, 16S GCN significantly differed across soil layers (F = 34.7, p < 0.001), sites (F = 10.3, p = 0.003), and their interaction (F = 4.6, p < 0.001). Across sites, the highest 16S GCN was found in Ol1 (mean = 2.34 ± 0.09), with Ol0 showing slightly lower values (2.29 ± 0.07, p = 0.049). 16S GCN then declined progressively through Olf (2.24 ± 0.08) and Ohf (2.16 ± 0.10), reaching the lowest values in Oh (2.08 ± 0.08, p < 0.001 vs Ol1). In the mineral soil, 16S GCN slightly increased again to 2.18 ± 0.07 in A10, which was significantly higher than Oh (p = 0.003), while Ohf and A5 did not differ significantly. At the site level, BBR consistently showed higher 16S GCN than KAN across all organic layers from Ol0 to Ohf (p < 0.05) and exceeded MIT in Ol1, Olf, and Ohf (p < 0.05) whereas MIT and KAN did not differ significantly with the exception of Ol0 and Oh, where MIT showed higher 16S GCN than KAN (p = 0.004 and p = 0.002, respectively). Significant site differences disappeared in the mineral soil layers. The full linear mixed model and post hoc test results are provided in Supplementary Tables S7 and S8.

### Site effects on the forest floor prokaryotes are layer-dependent

We calculated the prokaryotic core microbiome, defined as genera present across all sites within a given layer. While these core taxa represented only a small fraction of the total diversity (<10% across all layers), they accounted for a substantial proportion of the total read abundance. However, its abundance share varied across layers. The relative core microbiome abundance dropped sharply from the Ol0 (56.8%) to the Ol1 (23.6%). Subsequently, the core dominance slightly increased along the Olf and Ohf but showed a substantial increase in the Oh (77%) and mineral soil layers (∼70%) (Supplementary Table S9).

A repeated analysis, including only complete sites KAN and MIT, was performed to exclude biases in the Oh due to an unbalanced design. However, the same pattern was observed in the reduced dataset (Supplementary Table S10).

Furthermore, the prokaryotic core microbiome shifted significantly across the layers (Figure 4A). In the Ol0, the core community was dominated by *Hymenobacter, Sphingomonas*, and *Massilia*, which showed their highest relative abundances in this horizon. In the Ol1 layer, *Mucilaginibacter* and *Tardiphaga* appeared as characteristic core taxa. *Granulicella, Puia*, and *Burkholderia* were identified specifically in the Olf and Oh layers. *Acidothermus* and *Aquisphaera* peaked in the lower forest floor layers Ohf and Oh, with *Acidothermus* being the dominant core taxon across layers Ohf to A10. Finally, in the mineral soil layers, taxa such as *Candidatus Solibacter* and *Bryobacter* were part of the core prokaryotic community.

**Figure 4:**
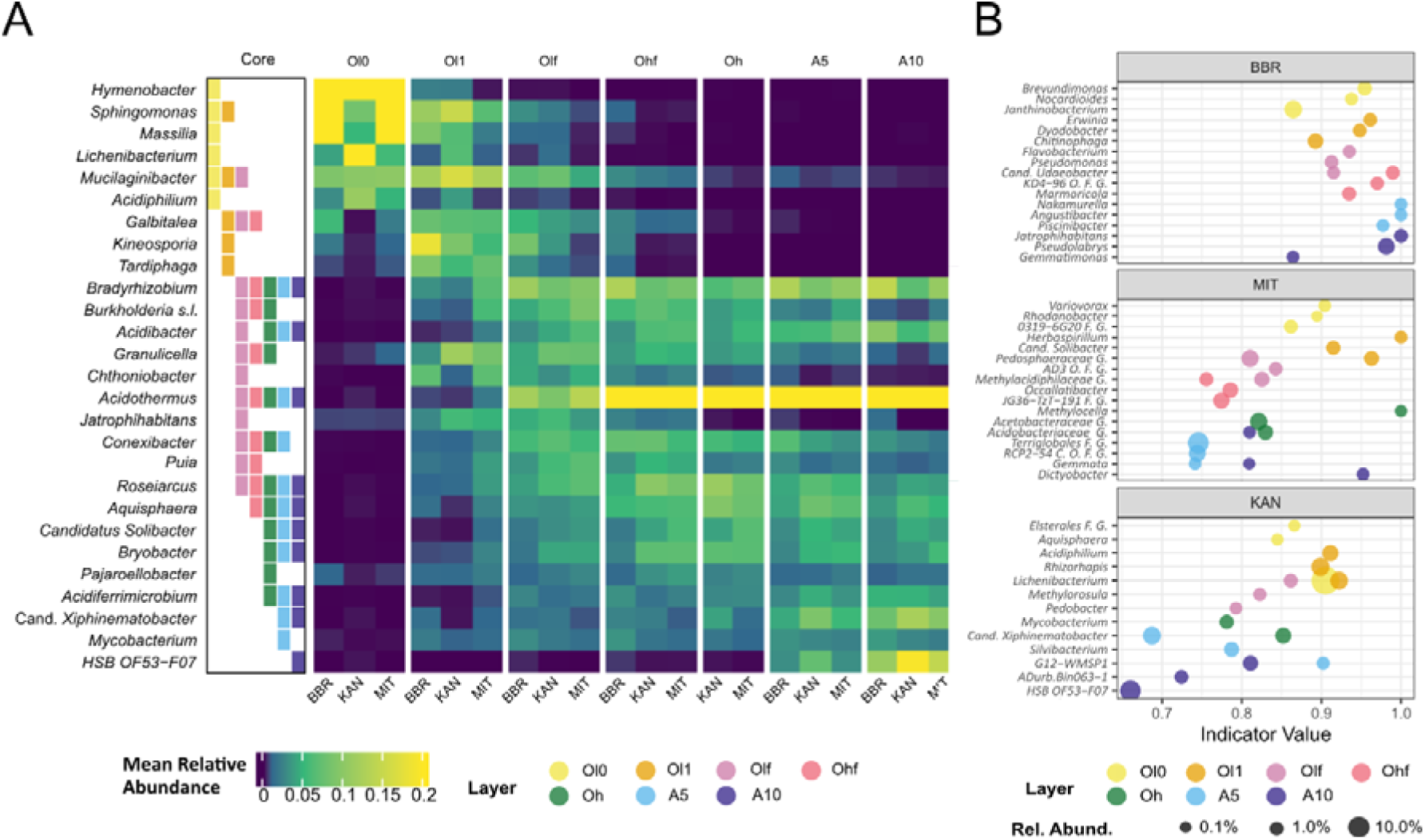
Site impact on the layer-specific forest soil prokaryotic community. (A) Shared prokaryotic genera (core microbiome) between the three surveyed sites for each layer of the forest floor. The core column indicates, in which FF layer the respective genus belongs to the core microbiome. (B) Indicator species analysis for each site and layer on genus level. Shown are the top three genera with the highest indicator value per layer and site.

Indicator species analysis was used to identify site-specific genera (Figure 4B), with the strongest indicator values occurring in different horizons depending on the site (OL layers at KAN and MIT, A layers at BBR). Accordingly, taxa associated with BBR exhibited the highest site specificity, reaching maximum indicator values in the mineral soil layers, which underscores the pronounced separation of the BBR communities in the NMDS analysis (compare Figure 2A). Key indicator genera for BBR included *Brevundimonas* (Ol0), *Erwinia* (Ol1), and *Candidatus Udaeobacter* in Olf and Ohf. *Nakamurella* and *Angustibacter* (A5), as well as *Jatrophihabitans* (A10), were identified as strong indicators for the mineral soil. In KAN, the genus *Lichenibacterium* was a consistent indicator for the upper forest floor layers (Ol0 to Olf), reaching a high relative abundance of 19.3% in the Ol0 horizon. Other prominent taxa included *Candidatus* Xiphinematobacter (Oh and A5) and G12-WMSP1 (A5 and A10). No significant indicator genera were identified for the Ohf layer. For MIT, *Variovorax* (Ol0), *Herbaspirillum* (Ol1), *Methylocella* (Oh), *Gemmata* (A5 and A10), and *Dictyobacter* (A10) were found as the strongest indicator genera. In contrast to BBR, indicator values of selected genera from KAN and MIT showed a declining trend with depth.

To evaluate how site-specific physicochemical parameters shape the forest soil prokaryotic community across the profile, we performed distance-based redundancy analysis (dbRDA) on weighted UniFrac distances (see full set of variables in Supplementary Table S11).

An overall dbRDA with forward selection of variables explained a large proportion of variance across the dataset, strongly reflecting the forest soil stratification (adj. R^2^ = 72.4%, p < 0.001, Supplementary Table S12 and Figure S6).

However, layer-wise dbRDA revealed that the environmental parameters driving community differences between sites varied distinctly across horizons (Table 3). The selected environmental parameters explained between 39.7% (Olf) and 64.2% (A10) of the compositional variation across all models (p < 0.01).

**Table 3:**
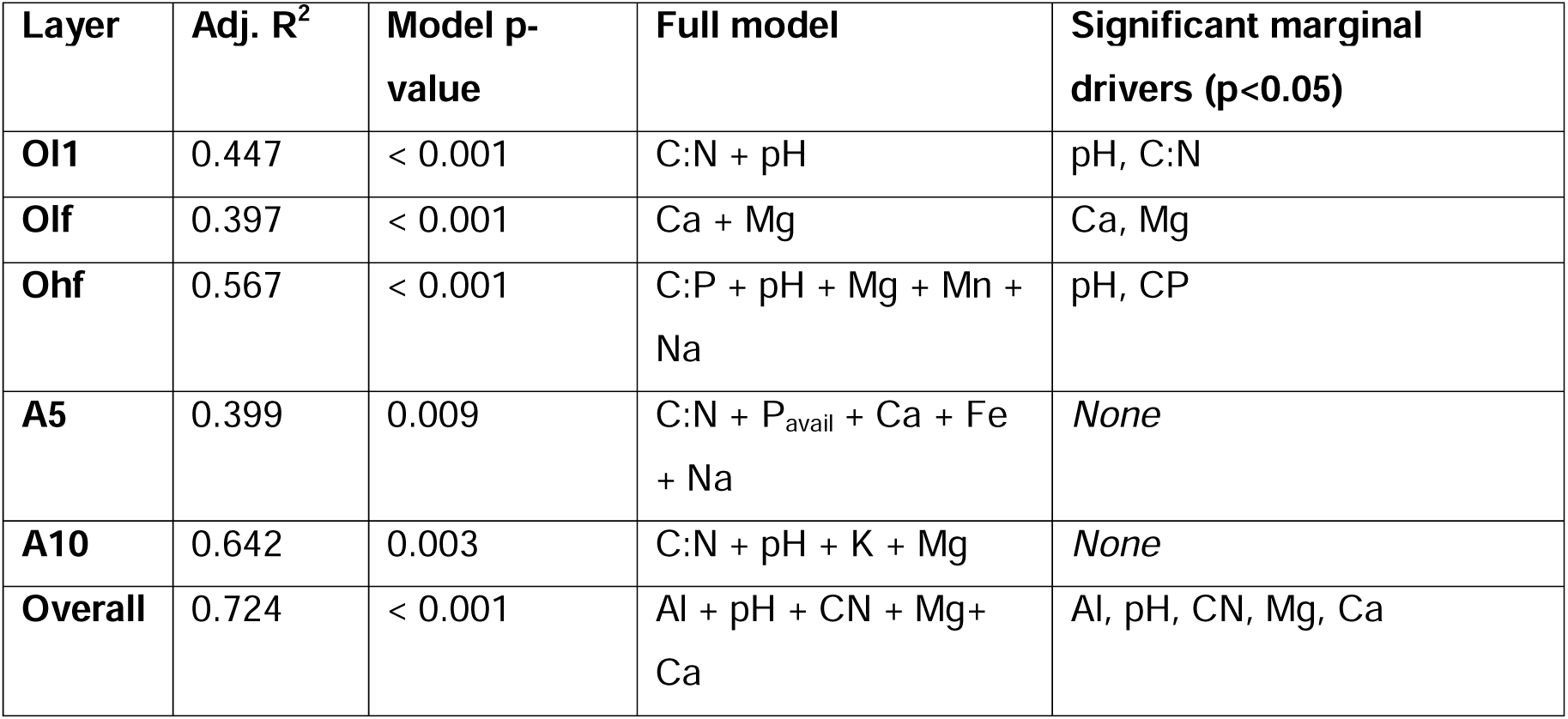
Environmental factors structuring prokaryotic community composition across forest soil layers. Environmental factors were selected in a multi-step process (see methods). The final predictors were then used for layer-wise distance-based RDA on weighted UniFrac distances to evaluate their explanatory power on the microbiome data. For the overall model, variables were selected by forward selection (ordiR2step). Significant marginal drivers were identified by subsequent permutational tests (999 permutations, p < 0.05).

In the litter layer Ol1, pH and C:N ratio were key drivers for community structure (p < 0.01), while exchangeable cations Ca and Mg were drivers in layer Olf (p < 0.05). In the Ohf layer, pH and C:P ratio were significant marginal drivers of community composition (p < 0.05). While the global models for the mineral soil layers A5 and A10 were significant and explained 39.9% and 64.2% of the variance, respectively, none of the marginal predictors was significant on its own. No significant model could be found for layer Oh. The full marginal test is provided in Supplementary Table S13.

## Discussion

### Site conditions drive taxa identity at the profile boundaries

Site-specific influence on the forest soil prokaryotic community varied both in strength and in its environmental drivers along the vertical profile.

Specifically, the prokaryotic community of the Ol1 was driven by pH and the C:N ratio of FF material, consistent with the findings, that litter quality controls early-stage microbial assembly (Asplund et al., 2019; Bray et al., 2012; Djukic et al., 2018; García-Palacios et al., 2016). In the Olf, site differences were driven by calcium and magnesium. Calcium is released during structural plant litter degradation and facilitates bacterial colonisation and decomposition, explaining its significance in this layer (Osono & Takeda, 2004; Prescott & Vesterdal, 2021; Shabtai et al., 2023). Similarly, magnesium was associated with faster decomposition rate, for example, by increasing bacterial beta-glucosidase activity, and may become a limiting nutrient in later decomposition stages through rapid leaching (Berg et al., 2021; Clarholm & Skyllberg, 2013; Giachetti & Vivanco, 2024). Interestingly, the C:P ratio was a significant predictor of the community site differences in the Ohf layer, suggesting that forest soil P status impacts the forest floor prokaryotic microbiome (Bergkemper et al., 2016; Lang et al., 2017). The lack of significant drivers in the Oh layer is likely caused by reduced statistical power due to a smaller sample size. Drivers of community variation in mineral soil lacked individual significance, potentially due to ecological collinearity among the physicochemical parameters defining site properties.

Our results suggest site specific factors control the prokaryotic community composition most strongly at the boundary layers of the surveyed profile. While the litter layer communities are strongly driven by litter quality, this impact diminishes with depth in the FF. The increase in the number and diversity of niches during decomposition may lead to higher local heterogeneity, as well as an overall community convergence between sites. Consequently, the communities of the Olf and Ohf layer are less dependent on site conditions, provided these layers remain intact. Site impacts return in the mineral soil, where decreased diversity, a resource-limited environment, and the increased impact of mineral soil parameters likely select for a site-specific community once again.

This effect was especially pronounced for site Bad Brückenau, where the prokaryotic community became more distinct from the other sites with depth. Compared to Kandel and Mitterfels, which are situated on paragneiss, Bad Brückenau is characterized by a basalt bedrock and contains higher concentrations of phosphorus and manganese. Fittingly, the most prominent indicator taxa identified at BBR in the mineral soil belong to the phylum *Actinomycetota*. Members of this phylum harbour key inorganic phosphorus solubilization genes, such as *gcd*, enriched in P-rich forest soils (Bergkemper et al., 2016; Ye et al., 2025) and their abundance was found to increase following phosphate rock addition in acidic soils (Bongoua-Devisme et al., 2024).

### The Stratification of the prokaryotic FF Microbiome Follows Common Principles Across Sites

Despite the apparent site impacts, stratification was the main driver of the FF prokaryotic community, exemplified also by the dominance of common taxa across sites. Vertical stratification trends in both alpha diversity patterns and community composition were consistent across the studied sites.

Alpha diversity of prokaryotes shifted strongly with depth in the forest soil profile. Overall, the FF exhibited a higher alpha diversity compared to the mineral topsoil, confirming previous research (Mundra et al., 2021; Qiao et al., 2025). High-resolution vertical profiling demonstrated that the high diversity of the FF is primarily an additive effect of diverging microbial communities across degradation stages. Therefore, FF stratification is the primary driver of increased biodiversity relative to mineral soil. As a consequence, climate-induced loss of FF layers is expected to directly reduce the alpha diversity of the FF as a whole.

Reports regarding which specific decomposition stage or forest floor layer supports the highest prokaryotic diversity appear contradictory. While the highest diversity in the forest floor profile was usually found in the litter layer (Baldrian et al., 2012; López-Mondéjar et al., 2015), time-resolved studies focusing on decomposition observed increasing diversity with progressing decomposition (Buresova et al., 2019; Gołębiewski et al., 2019; Purahong et al., 2016). In our results, alpha diversity peaked in the Olf and Ohf layers. This diversity hotspot might be driven by an ecotone-like effect where the overlap of litter and humified material provides niches for prokaryotes adapted to both the degradation of readily available and more recalcitrant resources (H. Wang et al., 2024).

In contrast, phylum-level richness consistently increased with depth in the soil profile. The lower FF layers and mineral soil present a chemically more heterogeneous environment that may provide niches for phylogenetically diverse taxa with distinct metabolic strategies, enabling diverse new phyla to establish (Maillard et al., 2024). Furthermore, compared to the highly dynamic FF, the mineral soil presents a more stable environment and can therefore serve as a reservoir where prokaryotic taxa accumulate over time (Lennon & Jones, 2011).

Alpha diversity patterns were mirrored by changes in prokaryotic community composition, revealing a clear successional stratification. The overall phylum abundance patterns were consistent across the three surveyed sites and align well with patterns observed previously in temperate deciduous forests (López-Mondéjar et al., 2015; Mundra et al., 2021).

*Pseudomonadota* and *Bacteroidota* generally decreased along the profile, and this trend was supported by the layer-specific core microbiome changes. Genera such as *Sphingomonas*, *Hymenobacter*, and *Massilia* prevailed in the Ol0 layer. These taxa are documented as inhabitants of the phyllosphere and early colonisers of litter, where traits such as UV- and cryoresistance likely provide a competitive advantage (Purahong et al., 2016; Sedláček et al., 2019; Smets et al., 2022). They were, however, quickly replaced by carbon degradation specialists like *Mucilaginibacter* and Tardiphaga (Hu et al., 2025; Khomutovska et al., 2024) in the Ol1, corroborating the notion that phyllosphere taxa play a limited role in decomposition (Tláskal et al., 2016).

In contrast, *Acidobacteriota* and *Chloroflexota* gained abundance with depth, exemplified in the enrichment of acidobacterial core genera Candidatus *Solibacter* and *Bryobacter* in the mineral soil. This trend aligns with their ecological characterisation as robust oligotrophs capable of persisting under nutrient- and carbon-limiting conditions as well as low oxygen availability (Dedysh, 2019; Fierer et al., 2007; Filip & Tesařová, 2005; Halamka et al., 2023; Speirs et al., 2019).

Notably, the abundance of *Actinomycetota* and *Planctomycetota* peaked in the FF layers Ohf and Oh. Specifically, *Acidothermus* and *Aquisphaera* were identified as core genera with peak relative abundances in the layers with high content of organic fine material. *Acidothermus* possesses extensive cellulolytic capabilities (Barabote et al., 2009; Brabcová et al., 2016), while *Aquisphaera* is associated with the degradation of lignin-rich residues (Tláskal et al., 2017), allowing both genera to access recalcitrant carbon pools. Furthermore, while upper litter layers favoured a copiotrophic lifestyle, likely due to the abundance of labile carbon sources (Mundra et al., 2021), the Ohf and Oh horizons exhibited the overall lowest 16S GCN values.

In FF layers dominated by organic fine matter, recalcitrant compounds like lignin, cellulose, and microbial necromass accumulate (Maillard et al., 2023; Šnajdr et al., 2011; B. Wang et al., 2021), leading to the formation of humic substances, the degradation of which is a complex multistep process (Kulikova & Perminova, 2021). In combination with our results, these findings suggest that those layers represent an environment that selects for oligotrophic bacteria, which are usually characterised by slow growth, high stress tolerance, and the capacity to co-utilize diverse low-concentration substrates (M. Zhu & Dai, 2024).

Taken together, the consistent stratification patterns across biogeochemically distinct sites suggest that litter type and quality are the primary drivers of the FF microbiome and therefore of decomposition processes (Bradford et al., 2016). Shared beech litter input largely overrides site-specific differences in shaping overall community assembly.

### Implications for Forest Ecosystem Functioning under Climate Change

The functional role of the FF prokaryotic community has direct implications for forest nutrition, as, for example, beech trees concentrate their roots in the topsoil, including the FF (Meier et al., 2018; Schenk & Jackson, 2002). However, organically bound nutrients within the humic layers of the FF are not directly available to plants. Bacteria and fungi mobilise these recalcitrant nutrient pools, and root exudation may enhance this process via priming effects (Yan et al., 2023). Beyond nutrient mobilisation, the formation of humic layers itself depends on microbial transformation processes and the accumulation of microbial necromass, making bacteria integral to both the formation and the functioning of the humic layers.

Climate change significantly impacts the forest ecosystems, including the forest soil microbiome (Baldrian et al., 2023; Seidl et al., 2017; Vacek et al., 2023). The FF is particularly sensitive to such disturbances, as defining processes like litter decomposition are strongly climate-dependent (Krishna & Mohan, 2017). Increasing temperatures accelerate microbial activity and carbon mineralisation, leading to C losses from the FF and potentially the reduction or even loss of FF layers and their associated specialised prokaryotic community (Hagedorn et al., 2019; Wellbrock et al., 2017). Furthermore, warmer winters may sustain microbial activity, contributing to year-round C losses in temperate forests (Baldrian et al., 2023). Conversely, prolonged drought may halt microbial decomposition, leading to the accumulation of plant-inaccessible humic substances (Guidi et al., 2022). The resulting shifts threaten the oligotrophic community in the humified layers. Because these taxa often mediate specialised functions such as the breakdown of recalcitrant organic matter, their loss might be especially critical due to a lack of functional redundancy, ultimately risking forest nutrition and long-term ecosystem stability.

## Conclusion

Forest floors play a crucial role in carbon cycling and nutrient turnover and thus are important for a functioning forest ecosystem. Our results demonstrate that vertical stratification of the forest floor is a primary and additive driver of overall forest-soil prokaryotic diversity and community structure. At the same time, site-specific factors shaped the prokaryotic community, particularly in the litter layer and mineral soil. Climate change may directly affect the forest soil diversity and functional redundancy if distinct FF layers are reduced or lost. In particular, the loss of oligotrophic niches in the lower FF layers dominated by recalcitrant organic matter may disrupt trees’ nutrient acquisition under nutrient poor conditions. As this study targeted the prokaryotic community, it does not capture fungal diversity, which is central to litter decomposition, nor cross-kingdom interactions. Shotgun metagenomics is therefore the necessary next step. Simultaneously covering fungal and bacterial diversity while extending the analysis from taxonomy to function is crucial for understanding the specific metabolic capabilities sensitive to climate and forest change and, therefore, for predicting their effects on the forest ecosystem.

## Acknowledgements

This work has been supported by the German Research Foundation (DFG) (Grant 457330647), as part of the Research Unit 5315.

Please refer to the Methods section for AI use declaration

## Author contributions

**Sebastian Bibinger**: Conceptualisation, Data curation, Formal analysis, Investigation (sampling, amplicon sequencing), Visualisation, Writing – original draft.

**Jörg Niederberger**: Data curation, Investigation (sampling, physicochemical measurements), Methodology, Project administration, Writing – review & editing.

**Friederike Lang**: Conceptualisation, Funding acquisition, Methodology, Project administration, Writing – review & editing.

**Michael Schloter**: Resources, Supervision, Writing – review & editing.

**Stefanie Schulz**: Conceptualisation, Funding acquisition, Resources, Supervision, Writing – original draft, Writing – review & editing.

## Data availability

The raw 16S amplicon sequencing data are available in the NCBI Sequence Read Archive (SRA) under accession no. PRJNA1454719. The R code used for processing, statistical analysis, and figure generation is available on GitHub (https://github.com/SebastianBibinger/FF_stratification)

## Bibliography

Adams, M. B., Kelly, C., Kabrick, J., & Schuler, J. (2019). Temperate forests and soils. In Global Change and Forest Soils (pp. 83–108). Elsevier. 10.1016/B978-0-444-63998-1.00006-9

Apprill, A., Mcnally, S., Parsons, R., & Weber, L. (2015). Minor revision to V4 region SSU rRNA 806R gene primer greatly increases detection of SAR11 bacterioplankton. Aquatic Microbial Ecology, 75(2), 129–137. 10.3354/AME01753

Asplund, J., Kauserud, H., Ohlson, M., & Nybakken, L. (2019). Spruce and beech as local determinants of forest fungal community structure in litter, humus and mineral soil. FEMS Microbiology Ecology, 95(2), 232. 10.1093/FEMSEC/FIY232

Baldrian, P. (2017). Forest microbiome: Diversity, complexity and dynamics. FEMS Microbiology Reviews, 41(2), 109–130. 10.1093/FEMSRE/FUW040

Baldrian, P., Kolařík, M., Štursová, M., Kopecký, J., Valášková, V., Větrovský, T., Žifčáková, L., Šnajdr, J., Rídl, J., Vlček, Č., & Voříšková, J. (2012). Active and total microbial communities in forest soil are largely different and highly stratified during decomposition. The ISME Journal, 6(2), 248. 10.1038/ISMEJ.2011.95

Baldrian, P., López-Mondéjar, R., & Kohout, P. (2023). Forest microbiome and global change. Nature Reviews Microbiology, 21(8), 487–501. 10.1038/s41579-023-00876-4

Bani, A., Pioli, S., Ventura, M., Panzacchi, P., Borruso, L., Tognetti, R., Tonon, G., & Brusetti, L. (2018). The role of microbial community in the decomposition of leaf litter and deadwood. Applied Soil Ecology, 126, 75–84. 10.1016/J.APSOIL.2018.02.017

Barabote, R. D., Xie, G., Leu, D. H., Normand, P., Necsulea, A., Daubin, V., Médigue, C., Adney, W. S., Xin, C. X., Lapidus, A., Parales, R. E., Detter, C., Pujic, P., Bruce, D., Lavire, C., Challacombe, J. F., Brettin, T. S., & Berry, A. M. (2009). Complete genome of the cellulolytic thermophile Acidothermus cellulolyticus 11B provides insights into its ecophysiological and evolutionary adaptations. Genome Research, 19(6), 1033. 10.1101/GR.084848.108

Barrett, M. (2022). ggokabeito: “Okabe-Ito” Scales for “ggplot2” and “ggraph” [Computer software]. https://malcolmbarrett.github.io/ggokabeito/

Bates, D., Mächler, M., Bolker, B., & Walker, S. (2015). Fitting Linear Mixed-Effects Models Using lme4. Journal of Statistical Software, 67(1), 1–48. 10.18637/jss.v067.i01

Berg, B., Sun, T., Johansson, M. B., Sanborn, P., Ni, X., Åkerblom, S., & Lönn, M. (2021). Magnesium dynamics in decomposing foliar litter – A synthesis. Geoderma, 382, 114756. 10.1016/J.GEODERMA.2020.114756

Bergkemper, F., Schöler, A., Engel, M., Lang, F., Krüger, J., Schloter, M., & Schulz, S. (2016). Phosphorus depletion in forest soils shapes bacterial communities towards phosphorus recycling systems. Environmental Microbiology, 18(6), 1988–2000. 10.1111/1462-2920.13188

Binkley, D., & Fisher, R. F. (2020). Ecology and management of forest soils. John Wiley & Sons, Inc.

Bongoua-Devisme, A. J., Kouakou, S. A. A., Kouadio, K.-K. H., & Lemonou Michael, B. F. (2024). Assessing the influence of diverse phosphorus sources on bacterial communities and the abundance of phosphorus cycle genes in acidic paddy soils. Frontiers in Microbiology, 15, 1409559. 10.3389/fmicb.2024.1409559

Brabcová, V., Nováková, M., Davidová, A., & Baldrian, P. (2016). Dead fungal mycelium in forest soil represents a decomposition hotspot and a habitat for a specific microbial community. New Phytologist, 210(4), 1369–1381. 10.1111/NPH.13849

Bradford, M. A., Berg, B., Maynard, D. S., Wieder, W. R., & Wood, S. A. (2016). Understanding the dominant controls on litter decomposition. Journal of Ecology, 104(1), 229–238. 10.1111/1365-2745.12507

Bray, S. R., Kitajima, K., & Mack, M. C. (2012). Temporal dynamics of microbial communities on decomposing leaf litter of 10 plant species in relation to decomposition rate. Soil Biology and Biochemistry, 49, 30–37. 10.1016/J.SOILBIO.2012.02.009

Buresova, A., Kopecky, J., Hrdinkova, V., Kamenik, Z., Omelka, M., & Sagova-Mareckova, M. (2019). Succession of microbial decomposers is determined by litter type, but site conditions drive decomposition rates. Applied and Environmental Microbiology, 85(24). 10.1128/AEM.01760-19

Callahan, B. J., McMurdie, P. J., Rosen, M. J., Han, A. W., Johnson, A. J. A., & Holmes, S. P. (2016). DADA2: High-resolution sample inference from Illumina amplicon data. Nature Methods, 13(7), 581–583. 10.1038/nmeth.3869

Clarholm, M., & Skyllberg, U. (2013). Translocation of metals by trees and fungi regulates pH, soil organic matter turnover and nitrogen availability in acidic forest soils. Soil Biology and Biochemistry, 63, 142–153. 10.1016/J.SOILBIO.2013.03.019

Currie, W. S. (1999). The responsive C and N biogeochemistry of the temperate forest floor. Trends in Ecology & Evolution, 14(8), 316–320. 10.1016/S0169-5347(99)01645-6

Davis, N. M., Proctor, D. M., Holmes, S. P., Relman, D. A., & Callahan, B. J. (2018). Simple statistical identification and removal of contaminant sequences in marker-gene and metagenomics data. Microbiome, 6(1), 226. 10.1186/s40168-018-0605-2

De Araujo Pereira, A. P., De Andrade, P. A. M., Bini, D., Durrer, A., Robin, A., Bouillet, J. P., Andreote, F. D., & Cardoso, E. J. B. N. (2017). Shifts in the bacterial community composition along deep soil profiles in monospecific and mixed stands of Eucalyptus grandis and Acacia mangium. PLOS ONE, 12(7), e0180371. 10.1371/JOURNAL.PONE.0180371

De Cáceres, M., & Legendre, P. (2009). Associations between species and groups of sites: Indices and statistical inference. Ecology, 90, 3566–3574. 10.1890/08-1823.1

Dedysh, S. N. (2019). Bryobacter. Bergey’s Manual of Systematics of Archaea and Bacteria, 1–5. 10.1002/9781118960608.GBM01667

Deutscher Wetterdienst. (2026). *Multi-*annual grids of annual mean air temperature and precipitation for Germany, 1991–2020 [Dataset]. https://opendata.dwd.de/climate_environment/CDC/grids_germany/annual/ (accessed 04/2026)

Djukic, I., Kepfer-Rojas, S., Schmidt, I. K., Larsen, K. S., Beier, C., Berg, B., Verheyen, K., Caliman, A., Paquette, A., Gutiérrez-Girón, A., Humber, A., Valdecantos, A., Petraglia, A., Alexander, H., Augustaitis, A., Saillard, A., Fernández, A. C. R., Sousa, A. I., Lillebø, A. I., … Tóth, Z. (2018). Early stage litter decomposition across biomes. Science of The Total Environment, *628–629*, 1369–1394. 10.1016/J.SCITOTENV.2018.01.012

Fierer, N., Bradford, M. A., & Jackson, R. B. (2007). Toward an ecological classification of soil bacteria. Ecology, 88(6), 1354–1364. 10.1890/05-1839

Filip, Z., & Tesařová, M. (2005). Microbial Processing of Humic Substances from Meadow and Forest Soils. Tree Species Effects on Soils: Implications for Global Change, 193–212. 10.1007/1-4020-3447-4_11

Floriancic, M. G., Allen, S., Meier, R., Truniger, L., Kirchner, J., & Molnar, P. (2022). Potential for significant precipitation cycling by forest-floor litter and deadwood. Authorea Preprints. 10.22541/AU.166209855.56534808/V3

Fox, J., & Weisberg, S. (2019). An {R} Companion to Applied Regression (Third). Sage. https://www.john-fox.ca/Companion/

Gao, G., Liu, Z., Wang, Y., Wang, S., Ju, C., & Gu, J. (2021). Tamm Review: Fine root biomass in the organic (O) horizon in forest ecosystems: Global patterns and controlling factors. Forest Ecology and Management, 491, 119208. 10.1016/J.FORECO.2021.119208

García-Palacios, P., Shaw, E. A., Wall, D. H., & Hättenschwiler, S. (2016). Temporal dynamics of biotic and abiotic drivers of litter decomposition. Ecology Letters, 19(5), 554–563. 10.1111/ELE.12590

Garnier, S., Ross, N., Rudis, R., Sciaini, M., Camargo, P. A., & Scherer, C. (2024). viridis(Lite)—Colorblind-Friendly Color Maps for R [Computer software]. 10.5281/zenodo.4679423

Giachetti, V. I., & Vivanco, L. (2024). Magnesium addition increases microbial metabolic efficiency during decomposition of Patagonian leaf litter. Plant and Soil, 507(1), 749–761. 10.1007/S11104-024-06766-9

Glassman, S. I., Weihe, C., Li, J., Albright, M. B. N., Looby, C. I., Martiny, A. C., Treseder, K. K., Allison, S. D., & Martiny, J. B. H. (2018). Decomposition responses to climate depend on microbial community composition. Proceedings of the National Academy of Sciences of the United States of America, 115(47), 11994–11999. 10.1073/PNAS.1811269115

Gołębiewski, M., Tarasek, A., Sikora, M., Deja-Sikora, E., Tretyn, A., & Niklińska, M. (2019). Rapid Microbial Community Changes During Initial Stages of Pine Litter Decomposition. Microbial Ecology, 77(1), 56–75. 10.1007/S00248-018-1209-X

Gu, Z. (2022). Complex heatmap visualization. iMeta, 1(3), e43. 10.1002/IMT2.43

Guidi, C., Frey, B., Brunner, I., Meusburger, K., Vogel, M. E., Chen, X., Stucky, T., Gwiazdowicz, D. J., Skubała, P., Bose, A. K., Schaub, M., Rigling, A., & Hagedorn, F. (2022). Soil fauna drives vertical redistribution of soil organic carbon in a long-term irrigated dry pine forest. Global Change Biology, 28(9), 3145–3160. 10.1111/gcb.16122

Gutachterausschuss Forstliche Analytik. (2014). Handbuch Forstliche Analytik (HFA). Bundesministerium für Ernährung und Landwirtschaft (BMEL).

Hagedorn, F., Gavazov, K., & Alexander, J. M. (2019). Above- And belowground linkages shape responses of mountain vegetation to climate change. Science, 365(6458), 1119–1123. 10.1126/science.aax4737

Halamka, T. A., Raberg, J. H., McFarlin, J. M., Younkin, A. D., Mulligan, C., Liu, X. L., & Kopf, S. H. (2023). Production of diverse brGDGTs by Acidobacterium Solibacter usitatus in response to temperature, pH, and O2 provides a culturing perspective on brGDGT proxies and biosynthesis. Geobiology, 21(1), 102–118. 10.1111/GBI.12525

Hao, J., Chai, Y. N., Lopes, L. D., Ordóñez, R. A., Wright, E. E., Archontoulis, S., & Schachtman, D. P. (2021). The Effects of Soil Depth on the Structure of Microbial Communities in Agricultural Soils in Iowa (United States). Applied and Environmental Microbiology, 87(4). 10.1128/AEM.02673-20

Hu, S., Wang, B. T., Li, T., Bu, S., Jin, C. Z., Jin, L., Ruan, H. H., Shin, K. S., & Jin, F. J. (2025). Successional patterns of microbial communities across various stages of leaf litter decomposition in poplar plantations. Frontiers in Microbiology, 16, 1628355. 10.3389/FMICB.2025.1628355

IUSS Working Group WRB. (2015). World Reference Base for Soil Resources 2014, 3rd edition, update 2015. International soil classification system for naming soils and creating legends for soil maps (World Soil Resources Reports No. 106). FAO.

IUSS Working Group WRB. (2022). World Reference Base for Soil Resources, 4th edition. International soil classification system for naming soils and creating legends for soil maps. International Union of Soil Sciences (IUSS).

Jiao, S., Chen, W., Wang, J., Du, N., Li, Q., & Wei, G. (2018). Soil microbiomes with distinct assemblies through vertical soil profiles drive the cycling of multiple nutrients in reforested ecosystems. Microbiome, 6(1), 1–13. 10.1186/S40168-018-0526-0

Kaiser, M., Ellerbrock, R. H., Wulf, M., Dultz, S., Hierath, C., & Sommer, M. (2012). The influence of mineral characteristics on organic matter content, composition, and stability of topsoils under long-term arable and forest land use. Journal of Geophysical Research: Biogeosciences, 117(G2), 2018. 10.1029/2011JG001712

Kassambara, A. (2026). ggpubr: “ggplot2” Based Publication Ready Plots [Computer software]. https://rpkgs.datanovia.com/ggpubr/

Khomutovska, N., Jasser, I., & Isidorov, V. A. (2024). Unraveling the Role of Bacteria in Nitrogen Cycling: Insights from Leaf Litter Decomposition in the Knyszyn Forest. Forests, 15(6), 1065. 10.3390/F15061065

Krishna, M. P., & Mohan, M. (2017). Litter decomposition in forest ecosystems: A review. *Energy*, Ecology and Environment, 2(4), 236–249. 10.1007/s40974-017-0064-9

Kulikova, N. A., & Perminova, I. V. (2021). Interactions between Humic Substances and Microorganisms and Their Implications for Nature-like Bioremediation Technologies. Molecules, 26(9), 2706. 10.3390/MOLECULES26092706

Kuznetsova, A., Brockhoff, P. B., & Christensen, R. H. B. (2017). lmerTest Package: Tests in Linear Mixed Effects Models. Journal of Statistical Software, 82(13), 1–26. 10.18637/jss.v082.i13

Laganière, J., Paré, D., & Bradley, R. L. (2010). How does a tree species influence litter decomposition? Separating the relative contribution of litter quality, litter mixing, and forest floor conditions. Canadian Journal of Forest Research, 40(3), 465–475. 10.1139/X09-208

Lahti, L., & Shetty, S. (2017). Tools for microbiome analysis in R (Version v.1.30.0) [Computer software]. https://github.com/microbiome/microbiome

Lang, F., Bauhus, J., Frossard, E., George, E., Kaiser, K., Kaupenjohann, M., Krüger, J., Matzner, E., Polle, A., Prietzel, J., Rennenberg, H., & Wellbrock, N. (2016). Phosphorus in forest ecosystems: New insights from an ecosystem nutrition perspective. Journal of Plant Nutrition and Soil Science, 179(2), 129–135. 10.1002/JPLN.201500541

Lang, F., Krüger, J., Amelung, W., Willbold, S., Frossard, E., Bünemann, E. K., Bauhus, J., Nitschke, R., Kandeler, E., Marhan, S., Schulz, S., Bergkemper, F., Schloter, M., Luster, J., Guggisberg, F., Kaiser, K., Mikutta, R., Guggenberger, G., Polle, A., … Chmara, I. (2017). Soil phosphorus supply controls P nutrition strategies of beech forest ecosystems in Central Europe. Biogeochemistry, 136(1), 5–29. 10.1007/S10533-017-0375-0

Lang, F., Prietzel, J., Hagedorn, F., Kaiser, K., Schulz, S., Mayer, M., Vesterdal, L., Bauhus, J., Bergmann, M., Bibinger, S., Bidartondo, M. I., Chen, J. O., De Jong, P., Doan, T. H., Haberstroh, S., Hahn, J., Hartmann, P., Khokon, A. M., Kohler, M., … Niederberger, J. (2025). Temperate forest floors: Ecosystem hub in transition? EarthArXiv. 10.31223/X5BB4G

Lavkulich, L. M. (1981). Methods manual: Pedology laboratory.

Lennon, J. T., & Jones, S. E. (2011). Microbial seed banks: The ecological and evolutionary implications of dormancy. Nature Reviews Microbiology, 9(2), 119–130. 10.1038/nrmicro2504

Lenth, R. V., Piaskowski, J., Banfai, B., Bolker, B., Buerkner, P., Giné-Vázquez, I., Hervé, M., Jung, M., Love, J., Miguez, F., Riebl, H., & Singmann, H. (2026). emmeans: Estimated Marginal Means, aka Least-Squares Means (Version 2.0.3) [Computer software]. https://cran.r-project.org/web/packages/emmeans/index.html

Lindahl, B. D., Ihrmark, K., Boberg, J., Trumbore, S. E., Högberg, P., Stenlid, J., & Finlay, R. D. (2007). Spatial separation of litter decomposition and mycorrhizal nitrogen uptake in a boreal forest. New Phytologist, 173(3), 611–620. 10.1111/J.1469-8137.2006.01936.X

Lladó, S., López-Mondéjar, R., & Baldrian, P. (2018). Drivers of microbial community structure in forest soils. Applied Microbiology and Biotechnology, 102(10), 4331–4338. 10.1007/S00253-018-8950-4

López-Mondéjar, R., Voříšková, J., Větrovský, T., & Baldrian, P. (2015). The bacterial community inhabiting temperate deciduous forests is vertically stratified and undergoes seasonal dynamics. Soil Biology and Biochemistry, 87, 43–50. 10.1016/J.SOILBIO.2015.04.008

López-Mondéjar, R., Zühlke, D., Becher, D., Riedel, K., & Baldrian, P. (2016). Cellulose and hemicellulose decomposition by forest soil bacteria proceeds by the action of structurally variable enzymatic systems. Scientific Reports, 6(1), 1–12. 10.1038/SREP25279

Maillard, F., Colin, Y., Viotti, C., Buée, M., Brunner, I., Brabcová, V., Kohout, P., Baldrian, P., & Kennedy, P. G. (2024). A cryptically diverse microbial community drives organic matter decomposition in forests. Applied Soil Ecology, 193, 105148. 10.1016/J.APSOIL.2023.105148

Maillard, F., Leduc, V., Viotti, C., Gill, A. L., Morin, E., Reichard, A., Ziegler-Devin, I., Zeller, B., & Buée, M. (2023). Fungal communities mediate but do not control leaf litter chemical transformation in a temperate oak forest. Plant and Soil, 489(1), 573–591. 10.1007/S11104-023-06040-4

Martin, M. (2011). Cutadapt removes adapter sequences from high-throughput sequencing reads. EMBnet.Journal, 17, 10. 10.14806/ej.17.1.200

McMurdie, P. J., & Holmes, S. (2013). phyloseq: An R package for reproducible interactive analysis and graphics of microbiome census data. PLoS ONE, 8(4), e61217. 10.1371/journal.pone.0061217

Meier, I. C., Knutzen, F., Eder, L. M., Müller-Haubold, H., Goebel, M.-O., Bachmann, J., Hertel, D., & Leuschner, C. (2018). The Deep Root System of Fagus sylvatica on Sandy Soil: Structure and Variation Across a Precipitation Gradient. Ecosystems, 21(2), 280–296. 10.1007/s10021-017-0148-6

Miao, J., Chen, T., Misir, M., & Lin, Y. (2024). Deep learning for predicting 16S rRNA gene copy number. Scientific Reports, 14(1), 1–14. 10.1038/S41598-024-64658-5

Mikryukov, V. (2025). metagMisc: Miscellaneous functions for metagenomic analysis [Computer software]. https://github.com/vmikk/metagMisc

Min, K., Zheng, T., Zhu, X., Bao, X., Lynch, L., & Liang, C. (2023). Bacterial community structure and assembly dynamics hinge on plant litter quality. FEMS Microbiology Ecology, 99(11), 1–10. 10.1093/FEMSEC/FIAD118

Minh, B. Q., Schmidt, H. A., Chernomor, O., Schrempf, D., Woodhams, M. D., Von Haeseler, A., Lanfear, R., & Teeling, E. (2020). IQ-TREE 2: New Models and Efficient Methods for Phylogenetic Inference in the Genomic Era. Molecular Biology and Evolution, 37, 1530–1534. 10.1093/molbev/msaa015

Mundra, S., Kjønaas, O. J., Morgado, L. N., Krabberød, A. K., Ransedokken, Y., & Kauserud, H. (2021). Soil depth matters: Shift in composition and inter-kingdom co-occurrence patterns of microorganisms in forest soils. FEMS Microbiology Ecology, 97(3), fiab022. 10.1093/FEMSEC/FIAB022

Naimi, B., Hamm, N. A. S., Groen, T. A., Skidmore, A. K., & Toxopeus, A. G. (2014). Where is positional uncertainty a problem for species distribution modelling. Ecography, 37, 191–203. 10.1111/j.1600-0587.2013.00205.x

Naylor, D., McClure, R., & Jansson, J. (2022). Trends in Microbial Community Composition and Function by Soil Depth. Microorganisms, 10(3), 540. 10.3390/MICROORGANISMS10030540

Oksanen, J., Simpson, G. L., Blanchet, F. G., Kindt, R., Legendre, P., Minchin, P. R., O’Hara, R. B., Solymos, P., Stevens, M. H. H., Szoecs, E., Wagner, H., Barbour, M., Bedward, M., Bolker, B., Borcard, D., Carvalho, G., Chirico, M., De Caceres, M., Durand, S., … Borman, T. (2025). vegan: Community Ecology Package [Computer software]. https://CRAN.R-project.org/package=vegan

Osono, T., & Takeda, H. (2004). Potassium, calcium, and magnesium dynamics during litter decomposition in a cool temperate forest. Journal of Forest Research, 9(1), 23–31. 10.1007/S10310-003-0047-X

Parada, A. E., Needham, D. M., & Fuhrman, J. A. (2016). Every base matters: Assessing small subunit rRNA primers for marine microbiomes with mock communities, time series and global field samples. Environmental Microbiology, 18(5), 1403–1414. 10.1111/1462-2920.13023

Ponge, J. F. (2013). Plant–soil feedbacks mediated by humus forms: A review. Soil Biology and Biochemistry, 57, 1048–1060. 10.1016/J.SOILBIO.2012.07.019

Posit team. (2025). RStudio: Integrated Development Environment for R. http://www.posit.co/

Prescott, C. E., & Vesterdal, L. (2021). Decomposition and transformations along the continuum from litter to soil organic matter in forest soils. Forest Ecology and Management, 498, 119522. 10.1016/J.FORECO.2021.119522

Purahong, W., Wubet, T., Lentendu, G., Schloter, M., Pecyna, M. J., Kapturska, D., Hofrichter, M., Krüger, D., & Buscot, F. (2016). Life in leaf litter: Novel insights into community dynamics of bacteria and fungi during litter decomposition. Molecular Ecology, 25, 4059–4074. 10.1111/mec.13739

Qiao, H., Zeng, Q., Martin, F., & Wang, Q. (2025). Impact of the soil layer on the soil microbial diversity and composition of Pinus yunnanensis at the Ailao Mountains subtropical forest. Frontiers in Microbiology, 16, 1558906. 10.3389/FMICB.2025.1558906

Quast, C., Pruesse, E., Yilmaz, P., Gerken, J., Schweer, T., Yarza, P., Peplies, J., & Glöckner, F. O. (2013). The SILVA ribosomal RNA gene database project: Improved data processing and web-based tools. Nucleic Acids Research, 41. 10.1093/nar/gks1219

R Core Team. (2025). R: A Language and Environment for Statistical Computing. https://www.R-project.org/

Raza, T., Qadir, M. F., Khan, K. S., Eash, N. S., Yousuf, M., Chatterjee, S., Manzoor, R., Rehman, S. ur, & Oetting, J. N. (2023). Unraveling the potential of microbes in decomposition of organic matter and release of carbon in the ecosystem. Journal of Environmental Management, 344, 118529. 10.1016/J.JENVMAN.2023.118529

Rowley, M. C., Grand, S., & Verrecchia, É. P. (2018). Calcium-mediated stabilisation of soil organic carbon. Biogeochemistry, 137, 27–49. 10.1007/s10533-017-0410-1

Sayer, E. J. (2006). Using experimental manipulation to assess the roles of leaf litter in the functioning of forest ecosystems. Biological Reviews, 81(1), 1–31. 10.1017/S1464793105006846

Sayer, E. J., Leitman, S. F., Wright, S. J., Rodtassana, C., Vincent, A. G., Bréchet, L. M., Castro, B., Lopez, O., Wallwork, A., & Tanner, E. V. J. (2024). Tropical forest above-ground productivity is maintained by nutrients cycled in litter. Journal of Ecology, 112(4), 690–700. 10.1111/1365-2745.14251

Schenk, H. J., & Jackson, R. B. (2002). Rooting depths, lateral root spreads and below-ground/above-ground allometries of plants in water-limited ecosystems. Journal of Ecology, 90(3), 480–494. 10.1046/j.1365-2745.2002.00682.x

Schliep, K. P. (2011). phangorn: Phylogenetic analysis in R. Bioinformatics, 27, 592–593. 10.1093/bioinformatics/btq706

Schroeter, S. A., Eveillard, D., Chaffron, S., Zoppi, J., Kampe, B., Lohmann, P., Jehmlich, N., von Bergen, M., Sanchez-Arcos, C., Pohnert, G., Taubert, M., Küsel, K., & Gleixner, G. (2022). Microbial community functioning during plant litter decomposition. Scientific Reports, 12(1), 1–10. 10.1038/s41598-022-11485-1

Sedláček, I., Pantůček, R., Králová, S., Mašlaňová, I., Holochová, P., Staňková, E., Vrbovská, V., Švec, P., & Busse, H. J. (2019). Hymenobacter amundsenii sp. Nov. Resistant to ultraviolet radiation, isolated from regoliths in Antarctica. Systematic and Applied Microbiology, 42(3), 284–290. 10.1016/J.SYAPM.2018.12.004

Seidl, R., Thom, D., Kautz, M., Martin-Benito, D., Peltoniemi, M., Vacchiano, G., Wild, J., Ascoli, D., Petr, M., Honkaniemi, J., Lexer, M. J., Trotsiuk, V., Mairota, P., Svoboda, M., Fabrika, M., Nagel, T. A., & Reyer, C. P. O. (2017). Forest disturbances under climate change. Nature Climate Change, 7(6), 395–402. 10.1038/nclimate3303

Shabtai, I. A., Wilhelm, R. C., Schweizer, S. A., Höschen, C., Buckley, D. H., & Lehmann, J. (2023). Calcium promotes persistent soil organic matter by altering microbial transformation of plant litter. Nature Communications, 14(1), 6609. 10.1038/s41467-023-42291-6

Smets, W., Spada, L. M., Gandolfi, I., Wuyts, K., Legein, M., Muyshondt, B., Samson, R., Franzetti, A., & Lebeer, S. (2022). Bacterial Succession and Community Dynamics of the Emerging Leaf Phyllosphere in Spring. Microbiology Spectrum, 10(2), e02420–21. 10.1128/SPECTRUM.02420-21

Šnajdr, J., Cajthaml, T., Valášková, V., Merhautová, V., Petránková, M., Spetz, P., Leppänen, K., & Baldrian, P. (2011). Transformation of Quercus petraea litter: Successive changes in litter chemistry are reflected in differential enzyme activity and changes in the microbial community composition. FEMS Microbiology Ecology. 10.1111/j.1574-6941.2010.00999.x

Šnajdr, J., Valášková, V., Merhautová, V., Cajthaml, T., & Baldrian, P. (2008). Activity and spatial distribution of lignocellulose-degrading enzymes during forest soil colonization by saprotrophic basidiomycetes. Enzyme and Microbial Technology, 43(2), 186–192. 10.1016/J.ENZMICTEC.2007.11.008

Speirs, L. B. M., Rice, D. T. F., Petrovski, S., & Seviour, R. J. (2019). The Phylogeny, Biodiversity, and Ecology of the Chloroflexi in Activated Sludge. Frontiers in Microbiology, 10, 468041. 10.3389/FMICB.2019.02015

Štursová, M., Žifčáková, L., Leigh, M. B., Burgess, R., & Baldrian, P. (2012). Cellulose utilization in forest litter and soil: Identification of bacterial and fungal decomposers. FEMS Microbiology Ecology, 80(3), 735–746. 10.1111/J.1574-6941.2012.01343.X

Tláskal, V., Voříšková, J., & Baldrian, P. (2016). Bacterial succession on decomposing leaf litter exhibits a specific occurrence pattern of cellulolytic taxa and potential decomposers of fungal mycelia. FEMS Microbiology Ecology, 92(11), fiw177. 10.1093/FEMSEC/FIW177

Tláskal, V., Zrůstová, P., Vrška, T., & Baldrian, P. (2017). Bacteria associated with decomposing dead wood in a natural temperate forest. FEMS Microbiology Ecology, 93(12), 157. 10.1093/FEMSEC/FIX157

Urbanová, M., Šnajdr, J., & Baldrian, P. (2015). Composition of fungal and bacterial communities in forest litter and soil is largely determined by dominant trees. Soil Biology and Biochemistry, 84, 53–64. 10.1016/J.SOILBIO.2015.02.011

Uroz, S., Ioannidis, P., Lengelle, J., Cébron, A., Morin, E., Buée, M., & Martin, F. (2013). Functional Assays and Metagenomic Analyses Reveals Differences between the Microbial Communities Inhabiting the Soil Horizons of a Norway Spruce Plantation. PLOS ONE, 8(2), e55929. 10.1371/JOURNAL.PONE.0055929

Vacek, Z., Vacek, S., & Cukor, J. (2023). European forests under global climate change: Review of tree growth processes, crises and management strategies. Journal of Environmental Management, 332, 117353. 10.1016/j.jenvman.2023.117353

Wachendorf, C., Frank, T., Broll, G., Beylich, A., & Milbert, G. (2023). A Concept for a Consolidated Humus Form Description—An Updated Version of German Humus Form Systematics. International Journal of Plant Biology, 14(3), 658–686. 10.3390/IJPB14030050

Wang, B., Liang, C., Yao, H., Yang, E., & An, S. (2021). The accumulation of microbial necromass carbon from litter to mineral soil and its contribution to soil organic carbon sequestration. CATENA, 207, 105622. 10.1016/J.CATENA.2021.105622

Wang, H., Wang, Z., Yu, J., Ma, C., Liu, L., Xu, D., & Zhang, J. (2024). The function and keystone microbiota in typical habitats under the influence of anthropogenic activities in Baiyangdian Lake. Environmental Research, 247, 118196. 10.1016/J.ENVRES.2024.118196

Wellbrock, N., Grüneberg, E., Riedel, T., & Polley, H. (2017). Carbon stocks in tree biomass and soils of German forests. Central European Forestry Journal, 63(2–3), 105–112. 10.1515/forj-2017-0013

Wickham, H. (2016). ggplot2: Elegant Graphics for Data Analysis. Springer-Verlag New York. https://ggplot2.tidyverse.org

Yan, S., Yin, L., Dijkstra, F. A., Wang, P., & Cheng, W. (2023). Priming effect on soil carbon decomposition by root exudate surrogates: A meta-analysis. Soil Biology and Biochemistry, 178, 108955. 10.1016/j.soilbio.2023.108955

Ye, D., Lin, Y., Liu, T., Zhang, X., Tang, Y., Wang, K., Huang, H., Yu, H., Wang, Y., He, X., & Li, T. (2025). Soil microbial community harboring key genes drives rhizosphere phosphorus mobilization of phosphorus-accumulating *Polygonum hydropiper*. Applied Soil Ecology, 213, 106314. 10.1016/j.apsoil.2025.106314

Zhang, X., Ni, X., Heděnec, P., Yue, K., Wei, X., Yang, J., & Wu, F. (2022). Litter facilitates plant development but restricts seedling establishment during vegetation regeneration. Functional Ecology, 36(12), 3134–3147. 10.1111/1365-2435.14200

Zhao, H., Zheng, W., Zhang, S., Gao, W., & Fan, Y. (2021). Soil microbial community variation with time and soil depth in Eurasian Steppe (Inner Mongolia, China). Annals of Microbiology, 71(1), 1–12. 10.1186/S13213-021-01633-9

Zhu, F., & Cheng, J. (2022). Comparison of the effects of litter decomposition process on soil erosion under simulated rainfall. Scientific Reports, 12(1), 1–13. 10.1038/S41598-022-25035-2

Zhu, J., Jiang, L., Chen, L., Jin, X., Xing, C., Liu, J., Yang, Y., & He, Z. (2023). Tree seedling growth allocation of Castanopsis kawakamii is determined by seed-relative positions. Frontiers in Plant Science, 14, 1099139. 10.3389/FPLS.2023.1099139

Zhu, M., & Dai, X. (2024). Shaping of microbial phenotypes by trade-offs. Nature Communications, 15(1), 4238. 10.1038/s41467-024-48591-9

