## Supplemental Figures for "Vertical stratification drives additive prokaryotic diversity in beech forest floors, while site conditions shape boundary layers"

Supplementary Figures

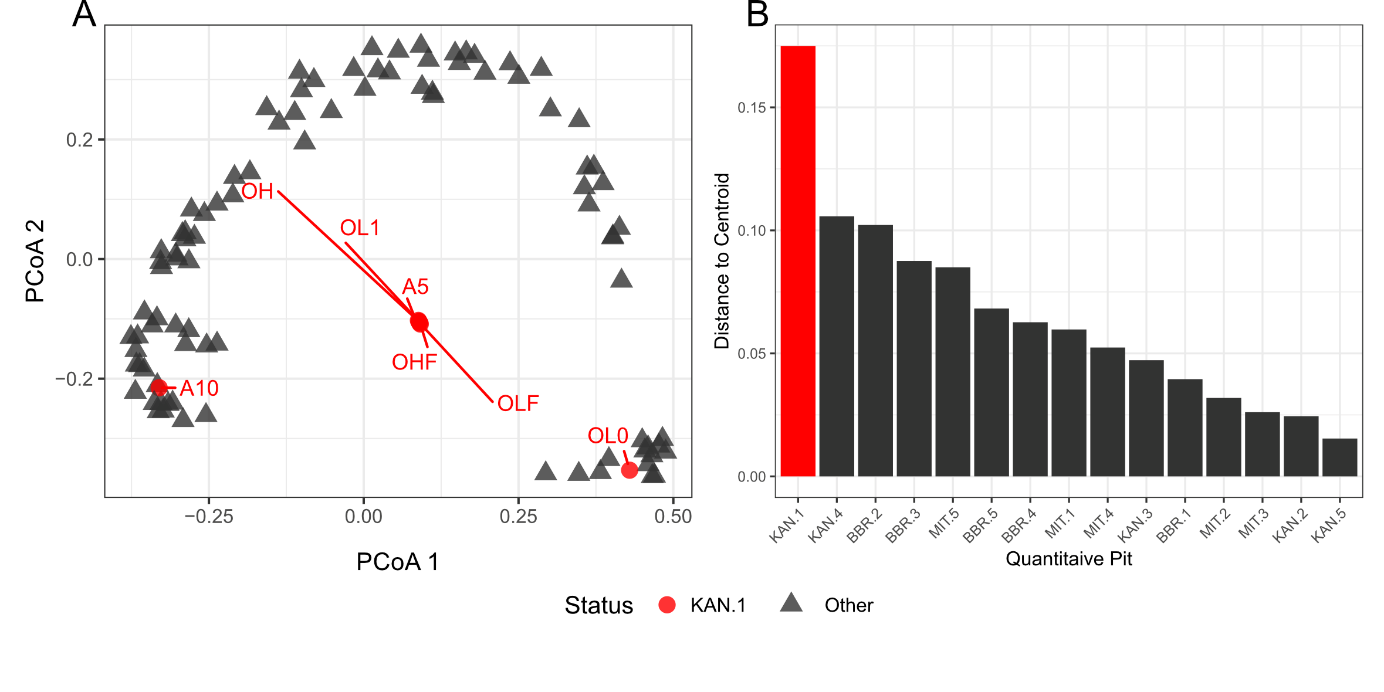
**Figure S1:** Identification of compositional outlier (Pit KAN-1). (A) Principal Coordinate Analysis (PCoA) of microbial communities based on Bray-Curtis dissimilarities on ASV level. Samples from q-pit KAN-are marked in red. (B) Euclidean distances of group centroids (q-pits) to the global dataset centroid. Q-pit KAN-1 exhibited a mean distance of 0.48, which is 1.65 times greater than that of the next most divergent q-pit, and was therefore excluded from further analysis.

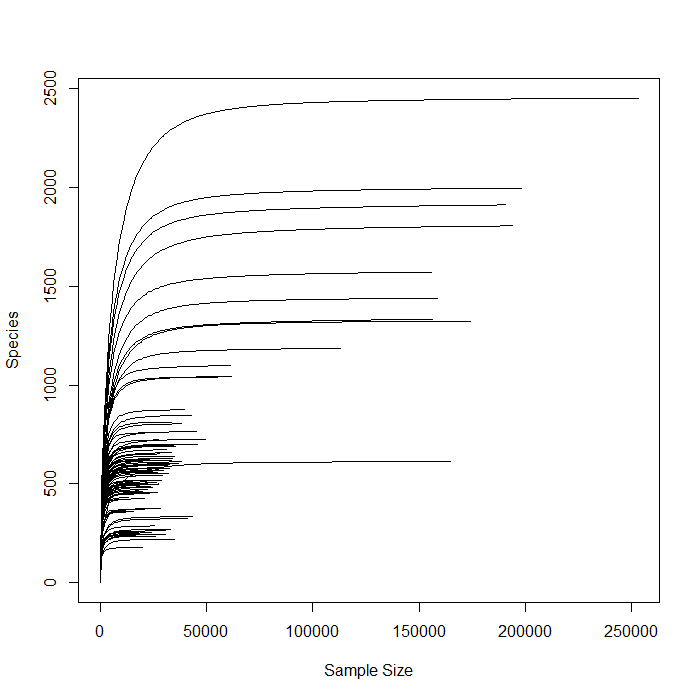

**Figure S2**: Rarefaction curves of observed amplicon sequence variant (ASV) richness across all samples, confirming sufficient sequencing depth for community-level analyses.

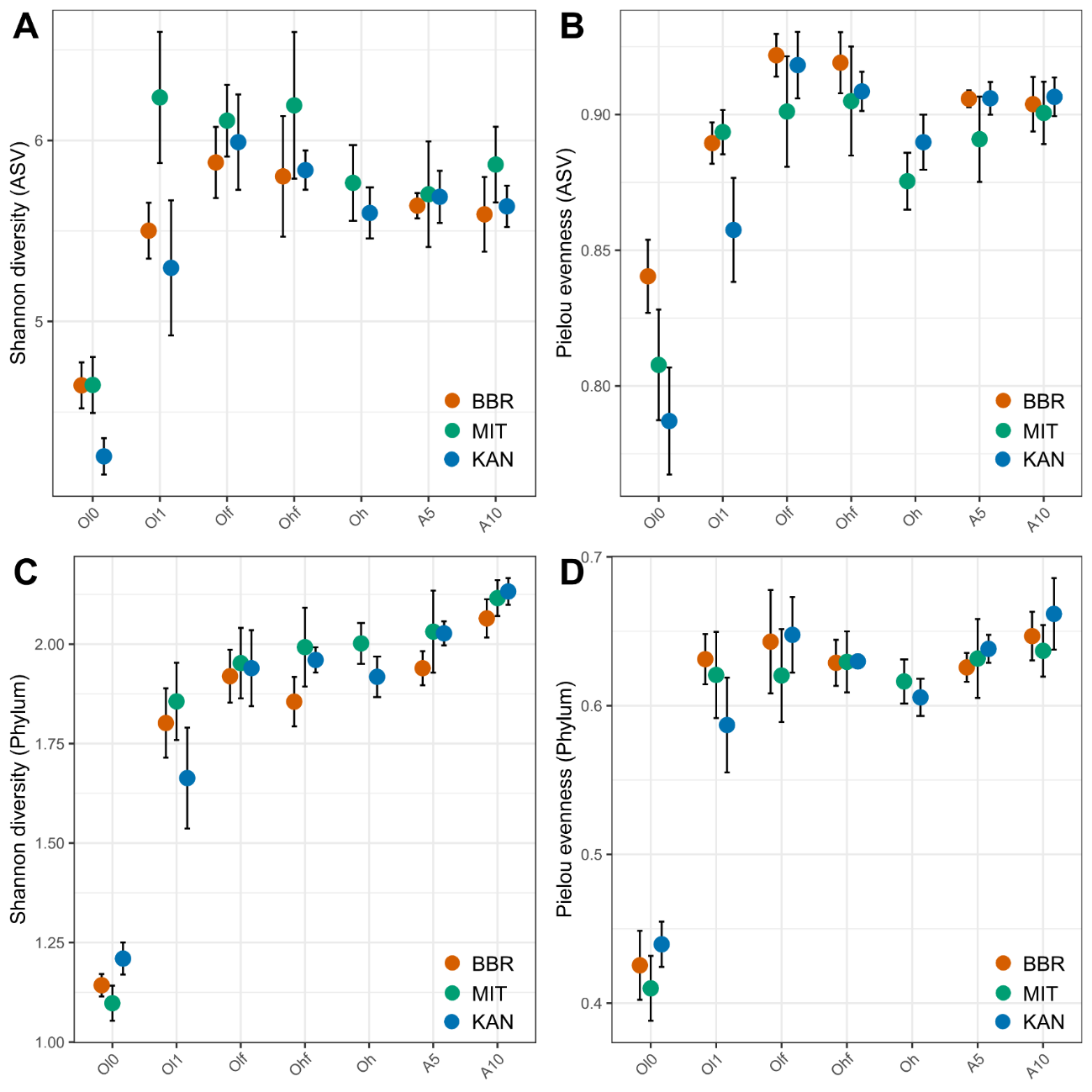

**Figure S3**: Alpha diversity along the forest floor profile at three German forest sites (Bad Brückenau, Mitterfels, Kandel). Shannon diversity (A, C) and Pielou's evenness (B, D) are shown at amplicon sequence variant (ASV) level (A, B) and phylum level (C, D). Indices were calculated on rarefied data (rarefaction depth = minimum sample size). Points represent the mean across replicate pits per site (n = 5 (BBR, MIT), n = 4 (KAN)); error bars indicate one standard deviation.

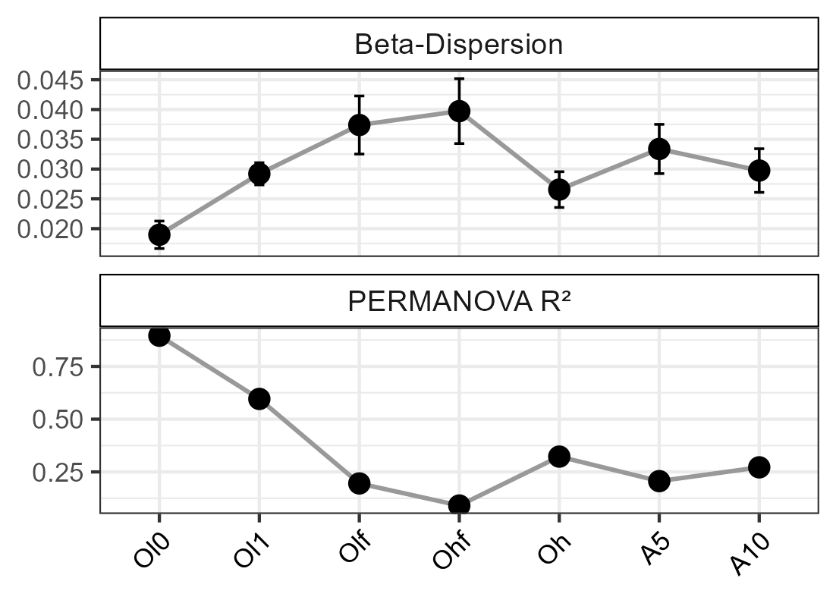

**Figure S4**: Beta-dispersion and PERMANOVA R² (site effect, based on Weighted UniFrac distances) of prokaryotic communities across forest floor layers at Kandel and Mitterfels. Analysis was performed on the reduced dataset excluding Bad Brückenau to avoid an unbalanced design due to the absence of layer Oh at Bad Brückenau.

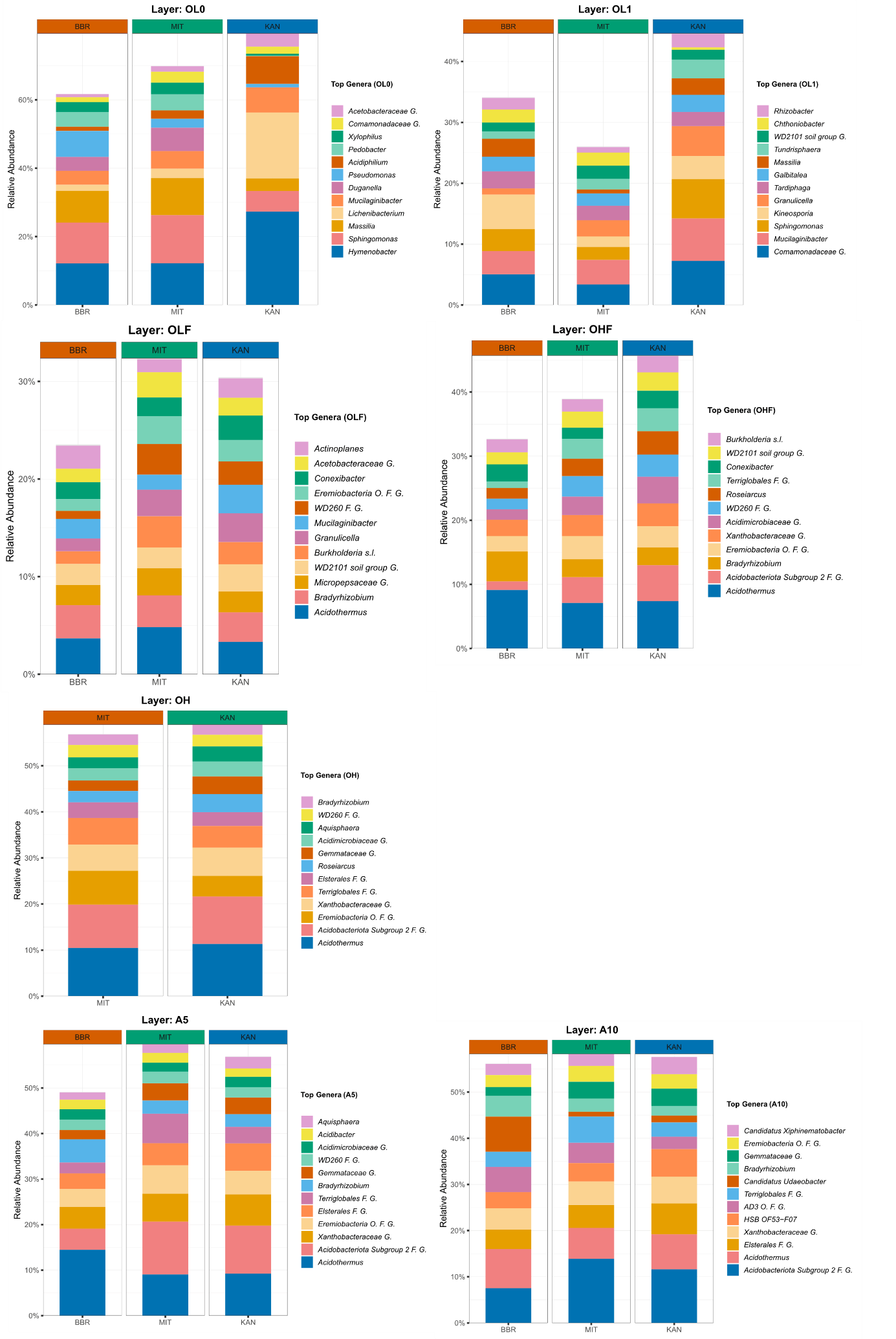

**Figure S5**: Relative abundance of the twelve most abundant genera across forest floor layers and mineral topsoil at the sampled beech forest sites Bad Brückenau (BBR), Mitterfels (MIT), and Kandel (KAN).

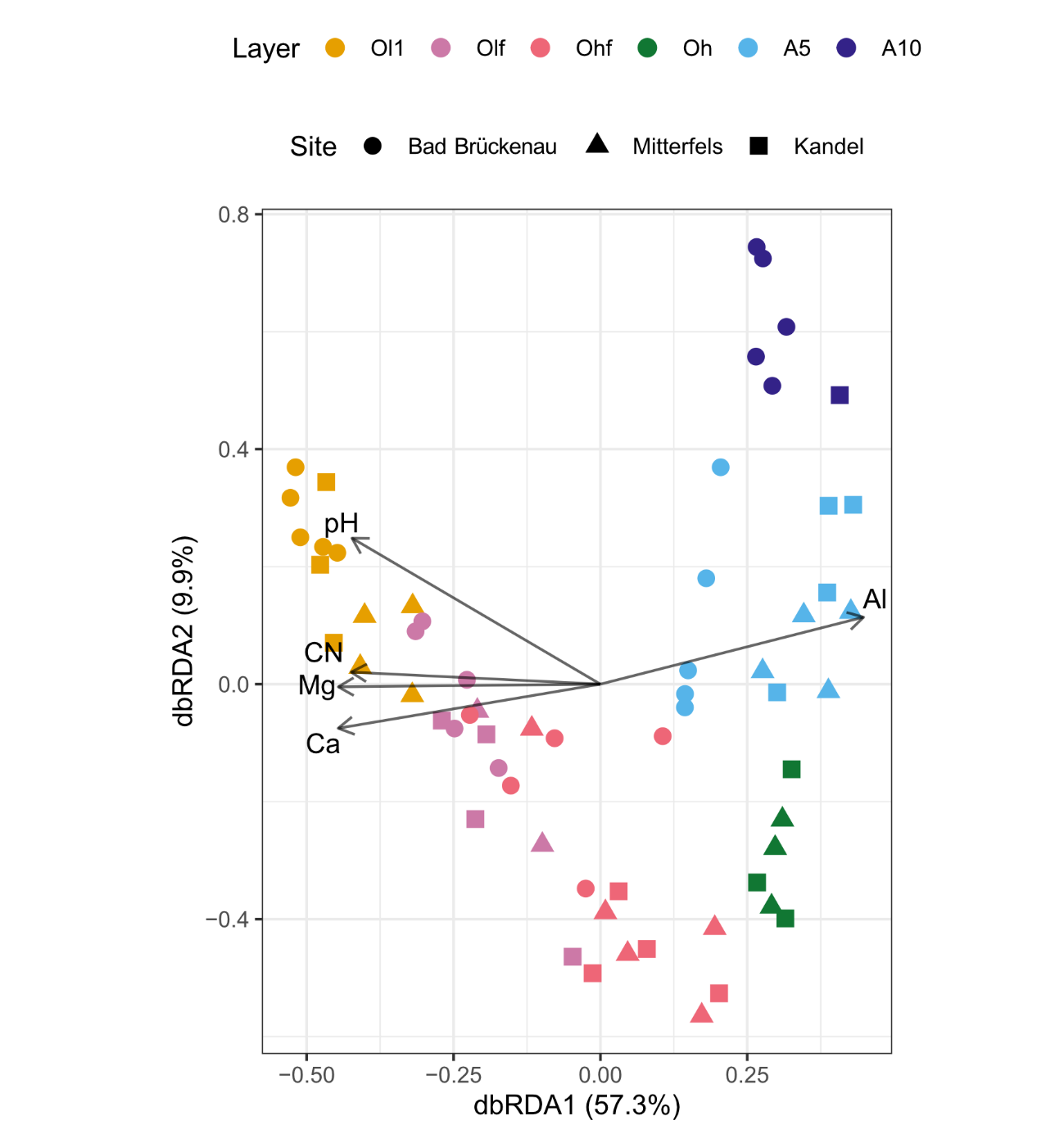

**Figure S6**: Global distance-based RDA model across all sites and layers. Environmental variables were selected through forward selection based on R^2^.
